# NLRP3 acts as a direct sensor of intracellular potassium ions

**DOI:** 10.64898/2026.03.12.707678

**Authors:** Ana Tapia-Abellán, Lukas Funk, Tobias Schäfer, Jelena Grga, Jane Torp, Corinna Gehring-Khav, Inga V. Hochheiser, Caroline Schönfeld, María Mateo-Tórtola, Fehime K. Eroglu, Gaopeng Li, Helmut Bischof, Robert Lukowski, Jasmin Kuemmerle-Deschner, Liudmila Andreeva, Christopher J. Farady, Matthias Geyer, Martin Frank, Alexander N.R. Weber

**Affiliations:** Institute of Immunology, Department of Innate Immunity, University of Tübingen, Auf der Morgenstelle 15, 72076 Tübingen, Germany; CMFI – Cluster of excellence (EXC 2124). “Controlling Microbes to Fight Infections”, Tübingen, Germany; iFIT – Cluster of excellence (EXC 2180). “Image-Guided and Functionally Instructed Tumor Therapies”, University of Tübingen, Germany; Institute of Structural Biology, University of Bonn, Venusberg-Campus 1, 53127 Bonn, Germany; Department of Internal Medicine VIII, University Hospital Tübingen, Röntgenweg 11, 72076 Tübingen, Germany; Pediatric Rheumatology and autoinflammation reference center Tübingen, Department of Pediatrics I, University Hospital Tübingen, Hoppe-Seyler-Str. 1, 72076 Tübingen, Germany; Experimental Pharmacology, Institute of Pharmacy, University of Tübingen, Auf der Morgenstelle 8, 72076 Tübingen, Germany; Novartis Biomedical Research, CH-4002 Basel, Switzerland; Biognos AB, Generatorsgatan 1, 41705 Göteborg, Sweden; German Cancer Consortium (DKTK), DKTK Partner Site Tübingen, 72076, Tübingen, Germany

**Keywords:** NLRP3, inflammasome, potassium, macrophage, inflammation

## Abstract

The NLRP3 inflammasome is a sentinel of cellular homeostasis, and its activation triggers the assembly of a molecular machinery that drives inflammation in infection, cardiovascular, metabolic, and neurodegenerative diseases. The majority of the many triggers known to activate NLRP3 are believed to induce potassium ion (K^+^) efflux from the cell as a fundamental danger signal for compromised cellular integrity. However, it has remained unclear how a reduction in intracellular K^+^ concentration is mechanistically translated into conformational changes in NLRP3 that promote inflammasome assembly, interleukin (IL)-1 release, and cell death. Here, we provide evidence that alterations in K^+^ levels directly regulate the conformation of the NLRP3 protein. In cell-free lysates derived from cell lines and primary blood immune cells high K^+^ concentrations stabilized a compact, protease-resistant structure resembling inhibitor-bound NLRP3, whereas low K^+^ conditions or the presence of a K^+^ chelator favored an open, more flexible and protease-accessible conformation. Notably, human NLRP3 remained responsive to K^+^ even when exogenously expressed in macrophage-like *Drosophila* cells or purified as recombinant protein. This indicates that K^+^ sensing occurs independently of cellular co-factors and is consistent with direct ion coordination. Of note, stimulation with the K^+^-independent NLRP3 agonist CL097 failed to recapitulate the conformational transition caused by K^+^ efflux inducer, nigericin. Moreover, pathogenic gain-of-function mutant variants of NLRP3 constitutively resembled the open and flexible protease-accessible conformation. Mapping K^+^-interactions by high-performance computation suggested that K^+^ ions populate the nucleotide binding pocket of the FISNA-NACHT module of individual NLRP3 chains but also stabilize face-to-face interactions within inactive oligomeric ‘cage’ assemblies via the NACHT-adjacent acidic loop. Collectively, our findings enable us to propose a mechanistic model of how intracellular K^+^ ions preclude NLRP3 activation prior to efflux and thus how NLRP3 responds to cellular danger as a direct K^+^ sensing protein.

## Introduction

Inflammation is a major cause of morbidity and mortality in humans. One major driver of innate-immune driven inflammation is NLRP3, a intracellular danger sensor that has been implicated in numerous inflammatory pathologies ranging from infections to cardiovascular and neurodegenerative disease (Weber, McManus et al. 2025). Whilst much of what we know about NLRP3 is based on cell and animal models, *NLRP3* gain-of-function mutations, which cause the autoinflammatory cryopyrin-associated periodic syndrome (CAPS), strikingly illustrate the potential of NLRP3 to cause systemic and local inflammation (Weber, McManus et al. 2025). In the absence of such auto-activating mutations, NLRP3 becomes active when cellular integrity is compromised upon infection (e.g. via pore-forming bacterial toxins such as nigericin or LukAB) or sterile insults, and, together with the adaptor ASC and an effector protease caspase-1, forms the inflammasome. This multi-protein complex processes substrates like pro-inflammatory IL-1 family cytokines for maturation and release, and gasdermin D for executing pyroptotic cell death (He, Wan et al. 2015, Liu, Zhang et al. 2016). In its inactive form, NLRP3 may be cytosolic, but a pool of NLRP3 resides in oligomeric form at the membranes of the trans-Golgi network (TGN) (Chen and Chen 2018, Andreeva, David et al. 2021) and is anchored to membrane lipids via a polybasic linker region encoded by exon 3 (Chen and Chen 2018, Bittner, Liu et al. 2021, Mateo-Tórtola, Hochheiser et al. 2023). Upon activation, oligomeric NLRP3 transits to the microtubule-organizing center (MTOC) where catalytically active inflammasomes are formed (Magupalli, Negro et al. 2020). On a structural level, TGN-bound inactive NLRP3 forms 10- or 12-membered oligomeric assemblies dubbed ‘cages’ (Andreeva, David et al. 2021, Hochheiser, Pilsl et al. 2022). The N-terminal PYD domain of each NLRP3 oligomer is supposed to be buried inside the inner cavity of the inactive ‘cage’ structures, in which the central NACHT and C-terminal LRR domains of inactive NLRP3 are interlocked by face-to-face (F2F) dimer contacts and assembled into a penta- or hexamer of dimers via back-to-back (B2B) contacts. Inhibitors like MCC950/CRID3 target the NACHT domain of NLRP3 and stabilize this inactive ‘closed’ conformation (Coll, Hill et al. 2019, Tapia-Abellan, Angosto-Bazarra et al. 2019, Dekker, Mattes et al. 2021, Hochheiser, Pilsl et al. 2022). dThe exact sequence of events is unknown, but the structure of an active NLRP3 ‘disk’ presents with a pivoted NACHT domain (Xiao, Magupalli et al. 2023, Yu, Matico et al. 2024). In this overall open conformation (Tapia-Abellan, Angosto-Bazarra et al. 2019), the exposed PYD domains nucleate the formation of ASC filaments (Lu, Magupalli et al. 2014, Hochheiser, Behrmann et al. 2022) that eventually engage active caspase-1 (Boucher, Monteleone et al. 2018). Whilst elegant structural and cell biological studies have provided informative snapshots of the three-dimensional conformations and subcellular locations of NLRP3 during activation, a central question in the field for decades has been how NLRP3-activating signals prompt conformational transition and inflammasome activation mechanistically.

Answering this question is a challenging endeavor due to the plethora of structurally and functionally disparate triggers of NLRP3. For example, ionophores and pore-forming toxins such as nigericin or LukAB (Perregaux and Gabel 1994, Mariathasan, Weiss et al. 2006, Liu, Pichulik et al. 2017), extracellular ATP acting via P2X7 ion channels (Perregaux and Gabel 1994, Mariathasan, Weiss et al. 2006), amyloid aggregates (Halle, Hornung et al. 2008, Masters, Dunne et al. 2010), crystals (Hornung, Bauernfeind et al. 2008, Duewell, Kono et al. 2010), compounds destabilizing lysosomes (Katsnelson, Rucker et al. 2015), and modulators of mitochondrial respiration (Gross, Mishra et al. 2016, Saller, Wohrle et al. 2025) all can activate NLRP3. The structural diversity of these agonists has given rise to the notion that NLRP3 is, firstly, a more general guardian of cellular integrity, and secondly, that agonist activity must somehow converge on a shared fundamental step that then acts upon NLRP3. Interestingly, pioneering work, even before the discovery of NLRP3, indicated that a change in the concentration of intracellular potassium ions (K^+^) could trigger IL-1β maturation (Perregaux and Gabel 1994). Later, the Tschopp group found that the drop in intracellular K^+^ from its high normal intracellular concentration of ∼150 mM down to below 90 mM could cause IL-1β maturation (Petrilli, Papin et al. 2007). As extracellular K^+^ concentration is low (∼3.5-5.0 mM), a steep transmembrane gradient is maintained by the activity of ion transporters, primarily the Na⁺/K⁺-ATPase (Palmer and Clegg 2016), and its disruption can lead to severe alterations in cellular homeostasis and to systemic dysfunction. Not surprisingly, a sudden drop in intracellular K⁺ ion concentration is cause for alarm, elicited via activation of the NLRP3 inflammasome. Although there clearly are K^+^-independent NLRP3 agonists (e.g. the mitochondria-active compounds imiquimod and CL097)(Gross, Mishra et al. 2016, Saller, Wohrle et al. 2025), K^+^ efflux as the common activation cue shared by multiple different NLRP3 agonists is considered a key paradigm in the inflammasome field (Munoz-Planillo, Kuffa et al. 2013).

However, given the importance of K^+^ for the integrity of NLRP3, there are surprisingly few studies investigating how the drop in K^+^ might orchestrate the switch from an inactive to an active conformation on a molecular level. Hypotheses have included a general disruption of ion homeostasis involving Ca^2+^ to trigger mitochondrial or ER stress; ROS and mitochondrial dysfunction; indirect sensing by other ancillary proteins; or direct sensing by NLRP3 (Tapia-Abellan, Angosto-Bazarra et al. 2021, Weber, McManus et al. 2025). As the effect of K^+^ has been evaluated mostly at the level of terminal pathway readouts (e.g. caspase-1 or IL-1β processing/release or cell death) rather than NLRP3 itself, none of these hypotheses to date has been sufficiently proven experimentally, leaving the most central tenet in the inflammasome field mechanistically unaddressed.

As an intracellular K^+^ sensing protein exists in bacteria (Ashraf, Josts et al. 2016), we sought to explore the simplest hypothesis, namely that NLRP3 directly senses the loss of K^+^. To this end, we took a reductionist approach, successively stripping overexpressed or endogenous NLRP3 from an intact mammalian cell environment down to purified proteins, monitoring the conformational impact of K^+^ in each setting. This showed that the conformation of NLRP3 is directly affected by K^+^ ion concentrations. Biochemical and high-performance computation using molecular dynamics simulation (MDS) enabled us to identify a critical interface in the NLRP3 FISNA-NACHT domain and the so-far enigmatic ‘acidic loop’ as key regions interacting directly with K^+^ ions and thus probably functioning as ion sensors and K^+^-stabilized conformational switches. These findings suggest a model in which K^+^ freezes NACHT flexibility and buffers the negative charges at the F2F interfaces of the NLRP3 oligomer, thereby maintaining a closed, inactive NLRP3 conformation. Conversely, K^+^ efflux, e.g., upon cell exposure to K^+^-dependent NLRP3 triggers, would relieve these ‘brakes’ and promote an open, active NLRP3 conformation. Consistently, K^+^-dependent conformational state transitions were observed in lysates derived from cells in which K^+^ efflux was triggered by nigericin, but not in cells treated with the K^+^-independent NLRP3 trigger CL097. Remarkably, hyperactive, disease-associated gain-of-function variants of NLRP3 proved to be insensitive to K^+^-mediated transition and constitutively exhibited an open conformation. Overall, our study not only provides the first direct experimental evidence that NLRP3 acts as a direct sensor that ‘guards’ intracellular K^+^ levels. It also integrates structural studies to provide a mechanistic model of how K^+^ efflux triggers a pivotal inflammatory response.

## Results

### Potassium ions modulate the conformation of human NLRP3 independently of an intact mammalian cellular environment

Although structural studies using cryo-EM and X-ray crystallography have greatly elucidated different conformational states of NLRP3 (Andreeva, David et al. 2021, Hochheiser, Pilsl et al. 2022, Xiao, Magupalli et al. 2023), they are currently not suited to interrogate NLRP3 conformation in living cells. Intramolecular bioluminescence resonance energy transfer (BRET) has been used successfully for monitoring overall structural changes of exogenously expressed NLRP3 in living cells, e.g. in response to both K^+^-dependent and -independent stimuli (Martin-Sanchez, Compan et al. 2016). However, as BRET solely measures the distance between the amino-terminal bioluminescent donor domain and the carboxyl-terminal fluorescent acceptor domain of overexpressed NLRP3, it is unable to define sub-structural conformational changes within NLRP3 domains, especially in untagged/endogenous NLRP3 and in human primary cells. Limited proteolysis is a low resolution but robust biochemical method for interrogating sub-structural conformational changes via protease accessibility (Fig. 1A) and was successfully applied to track binding of the inhibitor MCC950 to NLRP3 (Coll, Hill et al. 2019). To test whether this method could also report on sub-structural change induced by K^+^ in endogenously expressed NLRP3, lysates derived from LPS-primed THP-1 macrophages were prepared by sonication using two different buffers (see Methods for detailed composition): a high-K^+^ buffer containing 150 mM KCl buffer (hereafter referred to as K⁺ in figures) and a K^+^-free buffer in which KCl was replaced by 150 mM NaCl (hereafter referred to as Na⁺). Upon addition of pronase—a mixture of proteolytic enzymes from *Streptomyces griseus*—to lysates prepared in the K⁺-free, Na^+^-containing buffer, endogenous NLRP3 was completely degraded over time, as assessed by immunoblotting with an NLRP3-specific antibody (Fig. 1B). In contrast, in lysates prepared in the presence of K⁺ at a concentration comparable to intracellular levels (150 mM KCl), NLRP3 showed limited proteolysis, as verified by the increasing occurrence of a stable ∼50-kDa fragment (Fig. 1A, B). Notably, the overall proteolytic activity of pronase, as visualized through 2,2,2-trichloroethanol (TCE) gel incorporation throughout (e.g. Fig. S1A), was similar in the two lysate preparations, indicating that the stabilizing effect of K^+^ on endogenous NLRP3 was not due to lower enzymatic activity). Importantly, immunoblotting for another NLR protein of similar size, NLRC4 (also known as IPAF), a sensor of microbial patterns (Zhao, Yang et al. 2011), clearly showed that K^+^ specifically affects NLRP3 but not NLRC4 stability (Fig. 1B), which is in line with data showing that the NLRC4 inflammasome is K^+^-insensitive (Petrilli, Papin et al. 2007, Hwang, Park et al. 2012, Mangan, Akbal et al. 2024). Interestingly, the size of the K^+^-stabilized NLRP3 band was virtually identical to a band obtained by us and others upon pronase digestion of NLRP3 in the presence of the inhibitor MCC950 (Coll, Hill et al. 2019), under both K^+^-containing and K^+^-free conditions (Fig. 1C, quantified in Fig. 1D), suggesting that K^+^ ions also stabilize a closed, inactive conformation of NLRP3. Of note, K^+^ dose titrations in THP-1 lysates showed that ∼50 mM K^+^ were sufficient to stabilize the ∼50 kDa band (Fig. S1B, quantified in S1C), in agreement with published data showing that caspase-1 processing and later IL-1β release do not occur if K^+^ is maintained at ∼40 mM K^+^ or higher (Petrilli, Papin et al. 2007, Schmacke, O’Duill et al. 2022). That the changes in protease sensitivity were truly K^+^-dependent was further confirmed by the addition of poly-(sodium 4-styrene sulfonate) (PNaSS), a K^+^-specific chelator, which was able to switch NLRP3 back to the protease-accessible conformation, even in the 150 mM K^+^-containing lysate buffer (Fig. 1C, quantified in Fig. 1D). The stabilizing effect of K^+^ on NLRP3 protease sensitivity when quantified across experiments was statistically significant (Fig. 1D).

**Figure 1:**
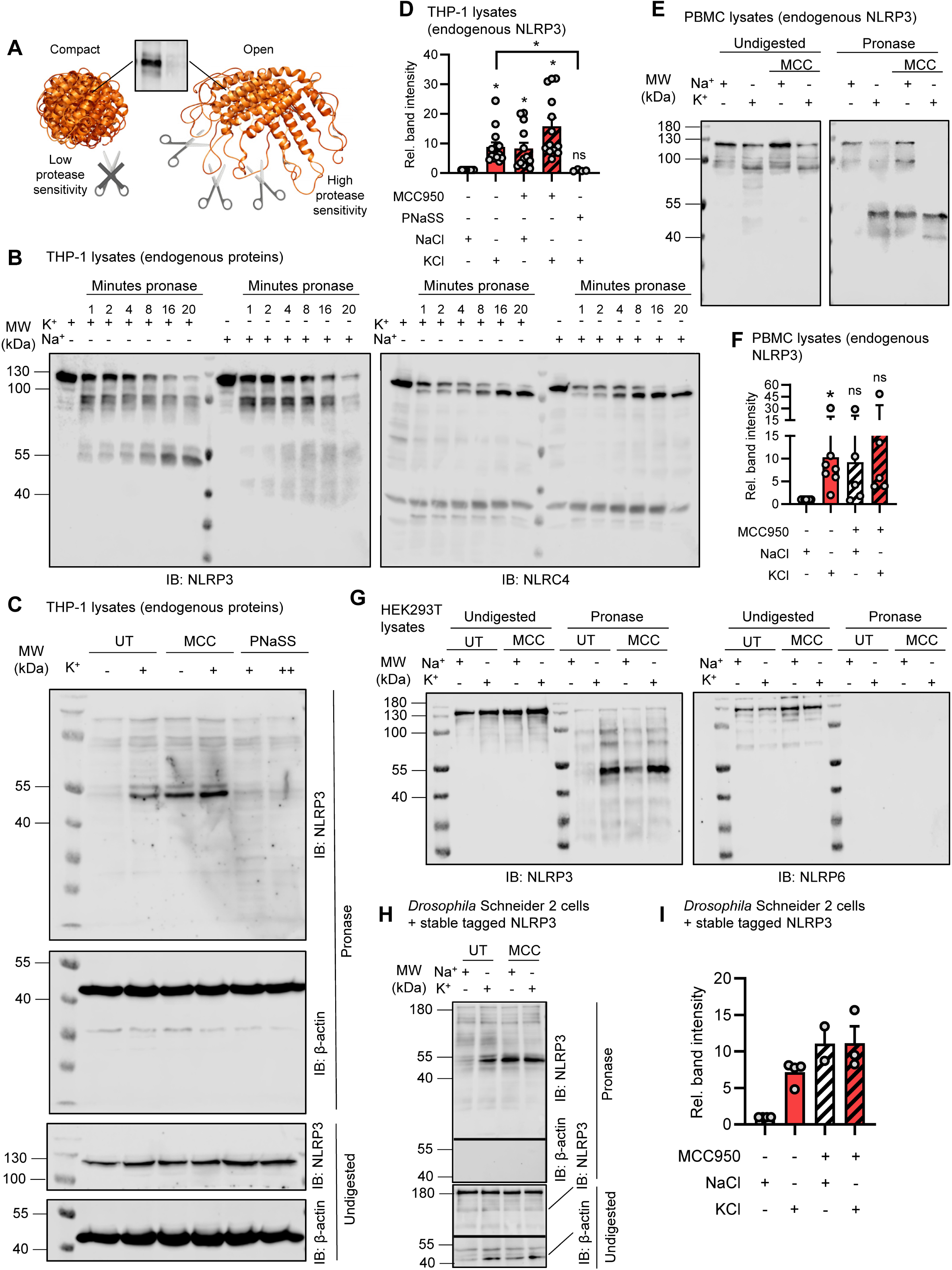
Potassium modulates endogenous NLRP3 conformational stability. **(A)** Principle of the detection of protein conformational states by pronase limited proteolysis assay. **(B)** Immunoblot of LPS-primed THP-1 cell lysates prepared by sonication in K^+^ (150 mM KCl) or Na^+^ (150 mM NaCl) buffer, followed by pronase addition for the indicated times. Anti-NLRP3 or anti-NLRC4 antibodies were used. Data are representative of n=3 independent experiments. **(C)** As in B but lysates were incubated ± MCC950 or ± PNaSS prior to pronase digestion, and anti-NLRP3 or anti-β-actin antibodies were used Data are representative of n=4-14 independent experiments. **(D)** Quantification of C showing combined data from n=4 independent experiments. One sample t-test compared to Na^+^ condition, ordinary one-way ANOVA comparing K^+^ and K^+^ with PNaSS. **(E)** As in C but with lysates from LPS-primed PBMC from healthy donor. Data are representative of n=5-7 different blood donors. **(F)** Quantification of E showing combined data from n=5-7 independent experiments (donors).One sample Wilcoxon signed rank test. **(G)** As in C but with lysates from HEK293T transiently transfected with YFP-NLRP3 or YFP-NLRP6. Data are representative of n=3 independent experiments. **(H)** As in C but with lysates from *Drosophila* S2 cells stably expressing human YFP-NLRP3-Luc. Data are representative of n=2-4 independent experiments. **(I)** Quantification of H showing combined data from n=2-4 independent experiments. one sample Wilcoxon signed rank test.

To investigate whether the conformational changes also occurred in a more physiological setting, we analyzed K^+^ sensitivity of endogenous NLRP3 in lysates derived from primary human immune cells, namely peripheral blood mononuclear cells (PBMCs). Pronase assays confirmed the same band protection as before (Fig. 1E, quantified in Fig. 1F). Collectively, these data show that adjusting K^+^ concentration in the lysis buffer alters the conformation of NLRP3, even endogenous NLRP3 from PBMC. We conclude that limited proteolysis of NLRP3 in cell lysates aligns well with sub-structural conformations in recent structural studies, namely an MCC950-stabilized inactive structure (Coll, Hill et al. 2019, Dekker, Mattes et al. 2021, Hochheiser, Pilsl et al. 2022) and that the method can report on NLRP3 conformation as a function of its environment’s K^+^ ion concentration: in lysates containing high K^+^ or MCC950, NLRP3 showed a compact structure (∼50 kDa band present), while the absence of K^+^ shifted NLRP3 to a more open, protease-accessible conformation (absence of the ∼50 kDa band). Limited proteolysis was therefore used in the following to investigate whether other factors are necessary for NLRP3 to respond to changes in K^+^ concentration or whether this is an intrinsic property of NLRP3 itself.

### K^+^-dependent conformational states are independent of inflammasome components or a mammalian cell context

The abovementioned experiments in cell lysates showed that K^+^-dependent conformational changes in endogenous NLRP3 were independent of a metabolically intact cellular environment or dynamic trafficking events. To investigate if other inflammasome components were required, we exogenously expressed YFP-NLRP3 or, as a control, another, K^+^-independent NLR protein of similar size, NLRP6 (Boegli, Bernard et al. 2026), in human embryonic kidney (HEK) 293T cells. This is a validated cellular model to study NLRP3 oligomerization and activation in the absence of the downstream inflammasome components ASC and caspase-1 (Bartok, Kampes et al. 2016). In this system, YFP-NLRP6 was equally degraded in K^+^ and Na^+^ conditions (Fig. 1G), whereas for YFP-NLRP3, only K^+^- or MCC950-containing lysates showed the protected band, suggesting critical downstream inflammasome components are not required for NLRP3-specific K^+^ sensitivity (Fig. 1G). However, other mammalian protein factors present in the lysate, e.g. SUGT1, HSP90 or NEK7, (Mayor, Martinon et al. 2007, He, Zeng et al. 2016, Schmid-Burgk, Chauhan et al. 2016) may still support or be required for conformational transition. To rule this out, we exogenously expressed human NLRP3, tagged with YFP and Luciferase, in macrophage-like *Drosophila melanogaster* Schneider 2 (S2) cells (Schneider 1972) and assayed lysates by limited proteolysis as before. Interestingly, almost identical results were obtained, i.e., the presence of a ∼50 kDa band in lysates containing 150 mM KCl and/or MCC950 (Fig. 1H, quantified in Fig. 1I) and its disappearance in lysates containing increasing concentrations of the K^+^ chelator, PNaSS (Fig. S1D and quantified in Fig. S1E). Comparable results were also observed when K^+^ was replaced by rubidium (RbCl) (Fig. S1F and quantified in Fig. S1G), suggesting that the presence of large, monovalent alkali metal cations with ≥3 electron shells is more relevant to NLRP3 conformation than any factors from a mammalian cellular environment.

### Recombinant purified NLRP3 undergoes a potassium-dependent conformational change

To conclusively demonstrate that NLRP3 itself responded to K^+^ concentration, we purified recombinant NLRP3 protein to homogeneity according to published protocols (Hochheiser, Pilsl et al. 2022). Of note, the very same full-length protein construct and purification procedure yielded decameric cages, both in the presence or absence of MCC950, as assessed by cryo-EM (Hochheiser, Pilsl et al. 2022). Interestingly, in the presence of high K^+^ and/or MCC950, limited proteolysis of highly pure NLRP3 yielded the same ∼50 kDa band as in cellular lysates (Fig. 2A, quantified in Fig. 2B), whereas pronase treatment of a protein A control protein, added to the same reaction prior to pronase addition, did not (Fig. S2A, quantified in Fig. S2B). This suggested not only that K^+^ stabilizes the same 3D conformation of purified NLRP3 as in cell lysates, but also that other cellular components – whether in THP-1, PBMCs, HEK293T or *Drosophila* S2 cells (*cf.* Fig. 1) – are not necessary to enable NLRP3 to adopt a more protease-resistant, compact conformation in response to K^+^. To validate this further, we conducted thermal stability assays, which measure an increase of a protein’s thermal stability as a result of ligand binding. This method was successfully used to demonstrate stabilization by ADP and/or inhibitors in non-oligomeric (Sharif et al, Nature, 2019) or truncated (i.e. only a FISNA-NACHT construct) NLRP3 proteins (Dekker, Mattes et al. 2021). The decameric NLRP3 showed a very high degree of thermal stability that could not be increased further by K^+^ addition, nor ADP or MCC950 (Fig. S2C), probably because this technique lacks the resolution to detect subtle structural changes in a large-sized, highly stable NLRP3 decamer in the absence of ASC. We therefore wondered whether NLRP3 K^+^ sensing was also related to its cage-like oligomeric conformation or represented an intrinsic property of the protein itself. To address this, we leveraged a functional NLRP3 variant lacking exon 3 (Δexon3) which does not form oligomers but assembles into smaller species such as dimers, both in vitro and when purified (Mateo-Tórtola, Hochheiser et al. 2023). Pronase assays using purified Δexon3 NLRP3 protein showed that this variant responded to K^+^, displaying the characteristic stable ∼50 kDa band (Fig. S2D, quantified in Fig. S2E). Moreover, K^+^-dependent band protection was also observable when Δexon3 NLRP3-mNeonGreen was stably reconstituted into *NLRP3* KO THP-1 cells (Mateo-Tórtola, Hochheiser et al. 2023) (Fig. S2F, quantified in S2G). The purified NLRP3 Δexon3 protein also underwent a significant increase in thermal stability with increasing K^+^ concentrations (Fig. S2H). Likewise, the presence of ADP led to an increase in stability, albeit not significantly (Fig. S2H).

**Figure 2:**
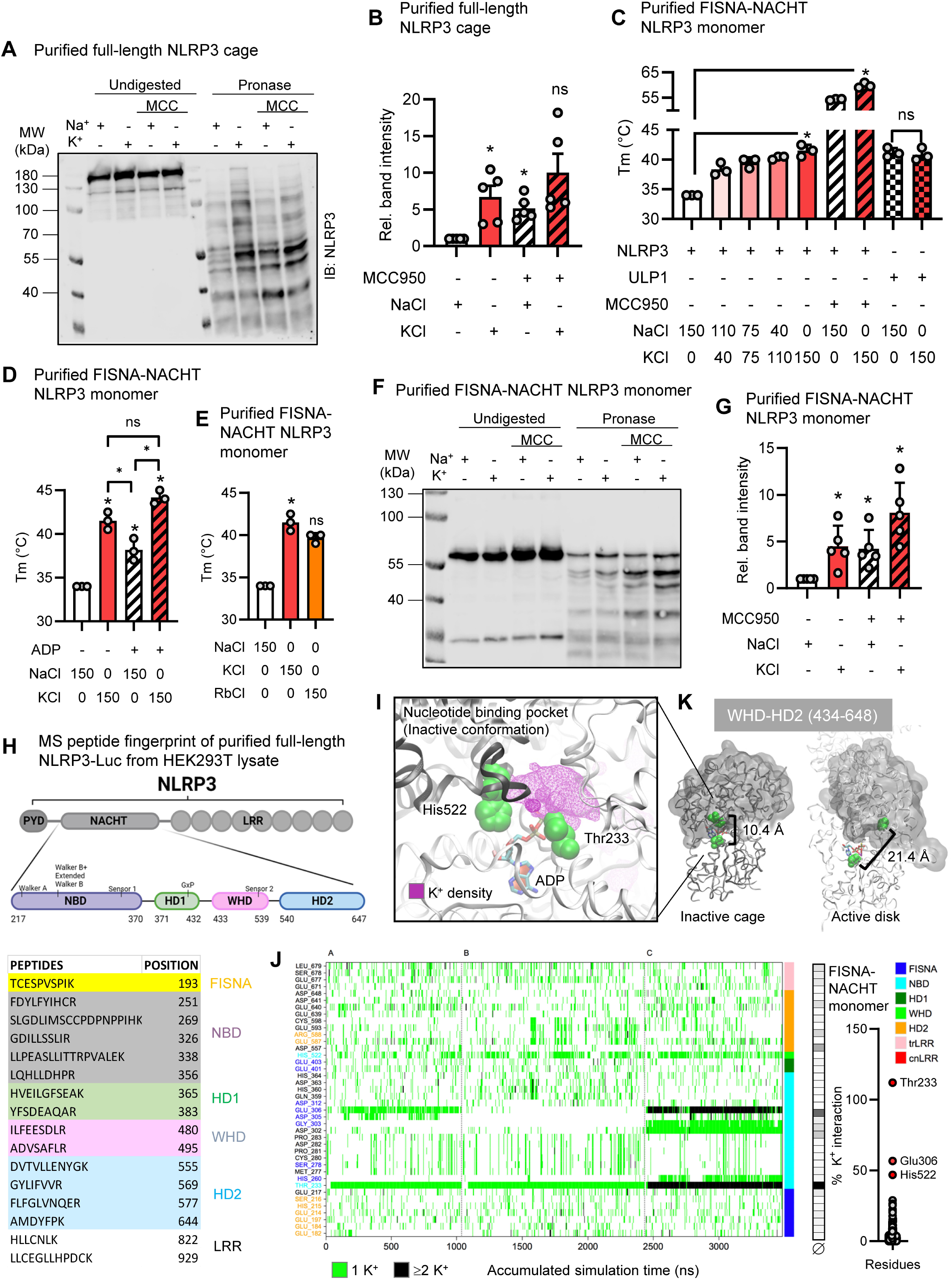
Potassium directly modulates conformation and stability of recombinant NLRP3. **(A)** Immunoblot of purified full-length NLRP3 (3-1036) in K^+^ (150 mM KCl) or Na^+^ (150 mM NaCl) buffer, with ± MCC950 added prior to pronase digestion. Anti-NLRP3 antibodies were used. Data are representative of n=5 independent experiments. **B)** Quantification of A showing combined data from n=5 independent experiments. One-way ANOVA with Dunnett’s multiple comparison test. **(C-E)** Melting temperature analysis (Tm, °C) of recombinant human FISNA-NACHT NLRP3 (131–679) or ULP1 in buffers containing increasing KCl and decreasing NaCl concentrations (total ion concentration 150 mM) ± MCC950 (C), or ± ADP (D) or using RbCl instead of KCl (E). Data show n=3 technical replicates (mean ±SD). One sample t-test. **(F)** As in A but with purified human FISNA-NACHT NLRP3 (131–679) protein. Data are representative of n=5 independent experiments. **(G)** Quantification of F combining n=5 independent experiments. One sample t-test. **(H)** Mass spectrometric peptide fingerprint of full-length NLRP3 immunoprecipitated with the specific NLRP3 antibody D4D8T from NLRP3-Luc-expressing HEK293T cells and purified after SDS-PAGE (Fig. S2I). Data from 1 independent experiment. **(I)** Representative snapshot from µs-scale MDS showing K⁺ ion density aggregated over time (magenta) within the nucleotide binding pocket of the NLRP3 NACHT domain and in proximity to Thr233. Trajectories and densities accumulated from n=3 independent MDS. **(J)** Time-resolved occupancy analysis of K⁺ ions within 5 Å sphere of individual NLRP3 residues (blue ID: residue is part of the NBD-HD1/WHD-HD2 interface; cyan ID: residue is within a 5 Å sphere of ADP, i.e. within nucleotide binding pocket; orange ID: residue is part of one of the NACHT domain surface loop region) across n=3 independent MDS (A-C) of the FISNA-NACHT structure, depicted in a heat map showing only residues with > 10% accumulated contact frequency (green = 1 K⁺ ion, black = ≥2 K⁺ ions; color code indicates to subdomains and grayscale heatmap indicates % occupancy averaged across all shown MDS). The graph depicts % occupancy for all FISNA-NACHT residues. **(K)** Comparison of NACHT subdomain orientations (NBD-HD1 part in white, WHD-HD2 in grey) in inactive cage structure (left) vs active disk (right). The distance between Thr233 and His522, both nucleotide binding pocket and K^+^ interacting residues, is taken as a proxy to illustrate the shift from inactive to active conformation.

To investigate whether the K^+^-stabilizing effect depended on the FISNA-NACHT domain (Tapia-Abellan, Angosto-Bazarra et al. 2021), and to rule out effects of NLRP3 multimerization applying to both full-length and Δexon3 NLRP3 proteins, we turned to a monomeric, purified FISNA-NACHT (131-679) protein construct with known structure (Dekker, Mattes et al. 2021). In thermal stability assays, titration of K^+^ led to a significant increase in protein stability, which could be further increased in the presence of MCC950 (Fig. 2C). Interestingly, the K^+^-dependent increase in melting temperature was concentration-dependent, with the strongest effect observable at the K^+^ concentration increase from 0 to 40 mM, and a more gradual increase at higher concentrations (Fig. 2C), whereas an established control protein, *Saccharomyces cerevisiae* ULP1 (Andreeva, David et al. 2021) was unaffected by K^+^ concentration. Addition of ADP also increased the thermal stability of the FISNA-NACHT construct in both NaCl and KCl buffers (Fig. 2D), and rubidium (RbCl) had a stabilizing effect on FISNA-NACHT very similar to K^+^ (KCl) (Fig. 2E). Similarly, in limited proteolysis experiments in the presence of high K^+^ concentration, the truncated monomeric FISNA-NACHT protein also yielded the ∼50 kDa band (Fig. 2F, quantified in Fig. 2G). These results show that the monomeric FISNA-NACHT construct of NLRP3 is able to directly, i.e., without the help of further co-factors, respond to K^+^ by adopting a protease-resistant compact structure, probably similar to the one published with MCC950 (Dekker, Mattes et al. 2021, Hochheiser, Pilsl et al. 2022). A more open, protease sensitive and less thermally stable conformation at low K^+^ concentrations is thus likely to correspond to a more active protein structure (Xiao, Magupalli et al. 2023), as would be expected upon K^+^ efflux in the wake of inflammasome triggering. Collectively, our data suggest that NLRP3 itself is a sensor of K^+^, which directly impacts its 3D structure.

### The NLRP3 FISNA-NACHT domain is conformationally affected by K^+^ sensing within its active site and the NBD-HD1/WHD-HD2 interface

Since the purified FISNA-NACHT was directly K^+^-sensitive, we sought to determine whether this domain corresponded to the observed stable ∼50 kDa band. We therefore expressed NLRP3-Luc in HEK293T cells, performed limited proteolysis experiments in the presence of 150 mM KCl, and analyzed the ∼50 kDa band, immunoprecipitated using an NLRP3 NACHT-specific antibody, on a Coomassie-stained gel (Fig. S2I), followed by mass spectrometric peptide fingerprinting. Evidently, the most N-terminal peptide mapped to the FISNA (from residue 193), but many additional peptides mapped to the NACHT domain (Fig. 2H), in good agreement with the immunoprecipitating NLRP3 NACHT antibody epitope surrounding Ala306 in the mouse NLRP3 NACHT. Although two peptides mapping to the LRR were also identified, these are likely contaminants as a polypeptide spanning from residue 193, in the most N-terminal peptide detected, to residue 822, in the closest detected LRR peptide, would have a predicted MW of ∼70 rather than ∼50 kDa. Altogether, this analysis, alongside the experiments with the purified FISNA-NACHT (*cf.* Fig. 2C-G), corroborated further that K^+^ acts directly on the FISNA-NACHT module, in good agreement with earlier cell-based studies (Tapia-Abellan, Angosto-Bazarra et al. 2021). This raised the question of how K^+^ ions may participate in activation at the level of the FISNA-NACHT domain.

Because unequivocal visualization of K^+^ ions requires exceedingly low resolution (<1.5 Å), we explored the plausibility and putative sites of K^+^ interactions within the FISNA-NACHT by molecular dynamics simulations (MDS), a computational method calculating atomic motions to model protein structure and dynamics over time based on classical physics force fields (Lee, Cheng et al. 2016, He, Man et al. 2020). Multiple parallel MDS (58,352 atoms in total, K^+^ ions corresponding to a concentration of 150 mM, microsecond (µs) timescale) were performed, and the interaction of K^+^ ions with individual residues was analyzed over time. This showed Thr233, a functionally relevant residue in the nucleotide binding pocket of NLRP3 (Brinkschulte, Fussholler et al. 2022), to reproducibly show the highest spontaneous binding and stable interaction with K^+^ ions, even on the µs timescale (Fig. 2I and 2J). Unexpected was the observation that K^+^ ions consistently populated the nucleotide binding pocket, which, in inactive NLRP3, is formed as a cleft between the NBD-HD1 and WHD-HD2 modules (Fig. 2I and 2K). Apart from Thr233, several other residues at this interface showed relatively high calculated K^+^ ion interactions, most notably Asp302, Glu306 and His522 (Fig. S2J). The latter appears coordinated, via a K^+^ ion, to Thr233 and ADP phosphate groups. In the active disk structure (Xiao, Magupalli et al. 2023), the WHD-HD2 is rotated by ∼90° around residue 437 (movement exemplified by depicting the distance between Thr233 and His522). Thus, we consider it likely that K^+^ contributes to the maintenance of an inactive NACHT conformation by stabilizing the NBD-HD1/WHD-HD2 interface. Collectively, our experimental and computational results indicate that the effects exerted by K^+^ ions involve stabilization of the core NACHT domain in its inactive conformation.

### K^+^ stabilization of the face-to-face dimer

Having shown experimentally that K^+^ directly affects the conformation of the FISNA-NACHT domain, we were interested to explore further how K^+^ might also affect the stability of the cage structure, i.e. multimer contacts, as ‘opening of the cage’ seems the critical checkpoint for NLRP3 to pass *en route* to activation. Based on (Mateo-Tórtola, Hochheiser et al. 2023) and the stability data for non-decameric Δexon3 (*cf.* Figs. S2D-H), we first explored NLRP3 dimer interfaces. Multiple parallel MDS runs (K^+^ ions corresponding to a molar concentration of 150 mM) were performed for either the F2F (165,108 atoms) or B2B (202,475 atoms) dimer assemblies derived from the human NLRP3 decamer (Hochheiser, Pilsl et al. 2022), and again used MDS to compute and map areas of K^+^ interaction and to calculate per-residue K^+^ dwell times (Figs. 3A, 3B and 3C). This not only confirmed the high K^+^ ion occupancy of Thr233 and the NBD-HD1/WHD-HD2 interface residues but, unexpectedly, also revealed pronounced occupancies in the F2F interface of the dimer involving (i) the acidic loop (Fig. 3B), a loop region between NACHT and LRR (Hochheiser, Pilsl et al. 2022), and (ii) the NLRP3 C-terminus (Fig. 3C). In addition, another site for K^+^ ions was found next to Asp648 and His674 (Fig. 3C). Conversely, the B2B interface, which is primarily hydrophobic in nature, showed practically no K^+^ ion occupancy (Fig. 3A).

**Figure 3:**
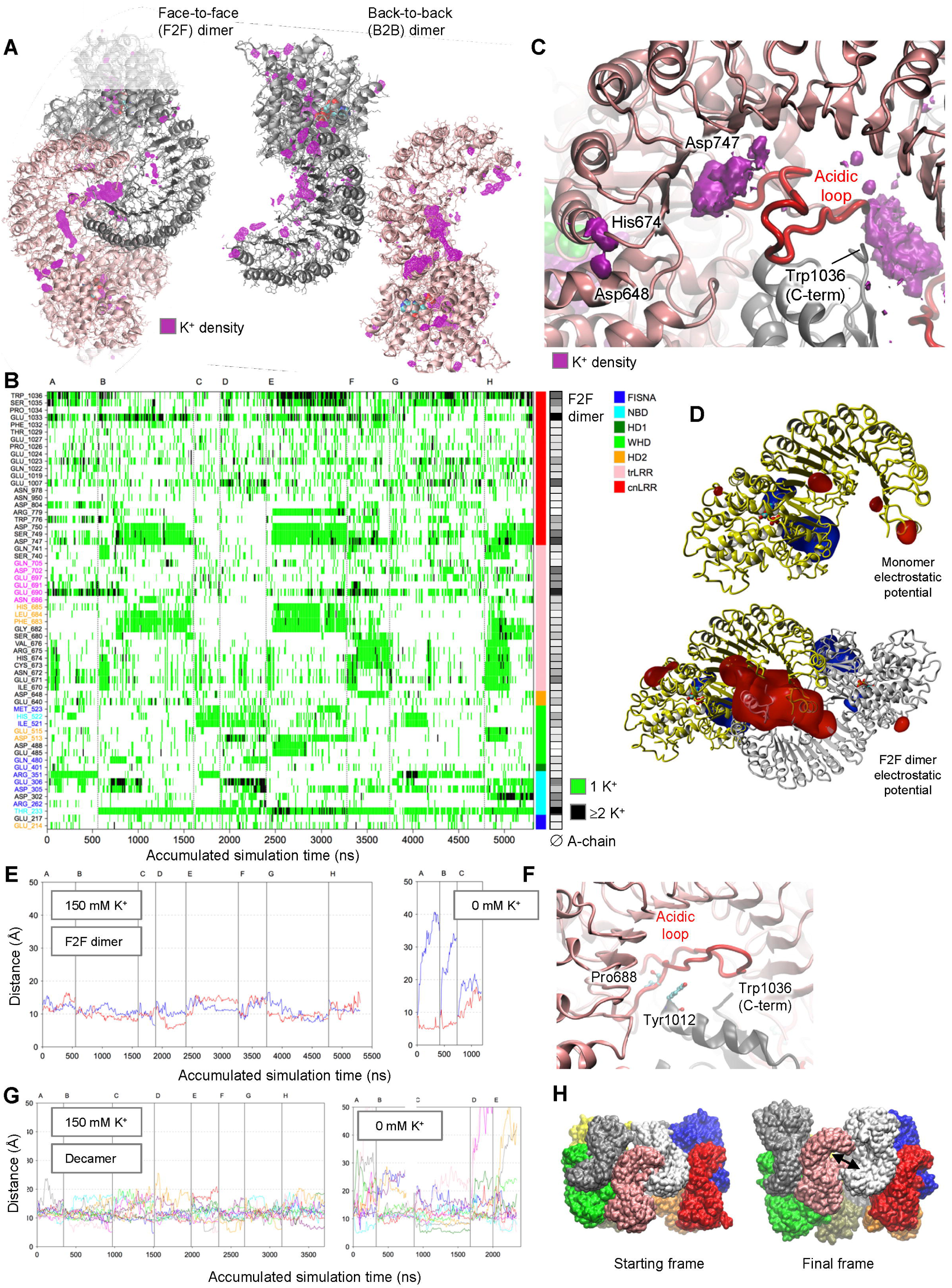
Potassium affects the NLRP3 NACHT domain and the F2F dimer interface in the oligomeric cage. **(A)** Structures of NLRP3 face-to-face (F2F, left) and back-to-back (B2B, right) dimers extracted from the decameric cage structure (PDB: 7PZC) and overlaid with K^+^ ion densities (magenta) calculated from respective molecular dynamics data. Data are representative from n=8 independent MDS. **(B)** Time-resolved occupancy analysis of K⁺ ions within 5 Å sphere of individual NLRP3 residues (pink ID: residue is part of the acidic loop (residues 686-706); blue ID: residue is part of the NBD-HD1/WHD-HD2 interface; cyan ID: residue is within a 5 Å sphere of ADP, i.e. within the nucleotide binding pocket; orange ID: residue is part of a NACHT domain surface loop region) across n=8 independent MDS (A-H) of the F2F dimer, depicted in a heat map showing only residues with > 10% accumulated contact frequency (green = 1 K⁺ ion, black = ≥2 K⁺ ions; color code indicates subdomains and grayscale heatmap indicates % occupancy averaged across all shown MDS for chain A of the F2F dimer). **(C)** F2F interface as in A but close-up on acidic loop (red) and highly K⁺-interacting residues **(D)** Isocontour plots of the electrostatic potential (ESP; negative = red, positive = blue) calculated using the Particle Mesh Ewald (PME) method for a monomeric NLRP3 (upper) and F2F dimer (lower) extracted from the 7PZC decamer cage structure. **(E)** Comparison of the stability of the F2F interface based on the distances between residues Pro688 and Tyr1012 (see F) calculated from μs MDS of the dimer. Trajectory of the two distances A:Pro688-B:Tyr1012 (blue) and A:Tyr1012-B:Pro688 (red) in n=8 independent MDS with 0.15 M KCl (letters A-H, left) and n=3 independent MDS without K^+^ ions (letters A-C, right). **(F)** Close up on acidic loop (red) and residues Pro688 and Tyr1012 as proxies for interface movement. **(G)** Comparison of the stability of the F2F interfaces based on the distances between residues Pro688 and Tyr1012 calculated from μs MDS of the decamer. n=8 independent MDS with 0.15 M KCl (letters A-H, left) and n=5 independent MDS without K^+^ ions (letters A-E, right). **(H)** Starting (left) and final (right) time frame of MDS ‘E’ from panel G (without K^+^ ions) of the complete NLRP3 decamer. One dissociated F2F interface is indicated with an arrow (*cf.* Movie S1).

We wondered about the electrostatic potential (ESP) present in the F2F interface and found a significant and widely distributed negative potential between the two LRR domains that strongly resembles the K^+^ ion density map obtained by MDS (Fig. 3D). In the NLRP3 monomer, only a greatly diminished negative potential was discernible in proximity to the acidic loop and the C-terminal end of the LRR domain, both of which come close upon F2F dimer formation. Therefore, we hypothesized that positive K^+^ ions could stabilize the F2F interaction by lowering the high negative ESP between the protein chains; consequently, K^+^ efflux would destabilize the F2F interface by eliminating this buffering mechanism. Indeed, MDS comparing the stability of F2F residue-residue contacts with and without K^+^ ions showed a drastic loss of stability and interface contacts in the absence of K^+^ ions, whereas most native residue–residue contacts in the F2F interface remained stable at 150 mM KCl (Fig. S3A). Moreover, the distance between residues Pro688 and Tyr1012 (Figs. 3E and 3F), used as a proxy for key interactions between the C-terminus of one LRR domain with the acidic loop of the other chain, fluctuated by only 5 Å (around the mean of 11 Å) over microseconds in the presence of K^+^ ions. Conversely, simulations without K^+^ ions over a similar timescale showed a nearly rapid and strong disintegration of the interface and separation of the LRRs (movement of >10 Å; example of last MDS frame provided in Fig. S3C).

Increasing the computational scale even further, we simulated the entire mega-Dalton (MDa) decameric cage structure of inactive NLRP3 (743,222 atoms in total; PDB entry 7pzc). Because of the size of the molecular system, an initial sampling on local computational infrastructure using AMBER forcefields (He, Man et al. 2020) was conducted over more than two months computer time but limited to 200 ns for each MDS. This nevertheless confirmed within this timescale a stabilizing effect of K^+^ ions on the F2F interfaces and relatively stable Pro688-Tyr1012 distances (Fig. S3D). In contrast, in simulations without K^+^ ions, the F2F interactions between two of the protein chains became instable within 100 ns simulation time and a complete dissociation of the subunits occurred (movement of >20 Å; Fig. S3D). Conversely, all B2B interfaces remained stable over the 200 ns simulation time also in the absence of K^+^ ions (Fig. S3D). In side-by-side analysis of MDS without K^+^, the fraction of native interface residue–residue contacts also remained stable for the B2B interface, whereas for the F2F it gradually declined (Fig. S3E).

To confirm these findings in even longer simulations of the decamer in a cubic box (892,158 atoms) and using a different force field (CHARMM, (Lee, Cheng et al. 2016)), 8 MDS of the NLRP3 cage in the presence of 150 mM KCl (accumulated timescale of >3.5 µs) and 5 MD simulations without K^+^ ions (accumulated >2 µs) were performed on a high-performance super-computing cluster. This fully reproduced the higher stability of the F2F interaction (approximated by the fraction of native residue–residue contacts, Fig. S3F) in the presence of K^+^, as well as the significant F2F conformational changes (estimated by the Pro688-Tyr1012 distance) and the disintegration of the interface in the absence of K^+^ (Fig. 3G, Movie S1 and Fig. 3H). As cage stability was found to be too high for experimental melting curve analysis (*cf.* Fig. S2C), we focused our experimental validation of these comprehensive computational data on the acidic loop as an essential integrator of K^+^ in the F2F interface. A construct with deletion of the acidic loop (Δac-loop; 42 residues, 683-724)) was designed, generated, expressed in HEK293T cells, and subjected to pronase degradation. In agreement with earlier data that a different acidic loop deletion can cause auto-activity (Hochheiser, Pilsl et al. 2022), a preliminary experiment showed that this Δac-loop NLRP3 variant did not show the band observed in WT NLRP3 upon K^+^ protection (Fig. S4A), indicating that it may be unable to adopt the stable, protease-resistant K^+^-bound conformation as predicted. Collectively, we provide computational and wet-lab data that suggest that K^+^ stabilization works at the level of both FISNA-NACHT and F2F dimer interface of decameric cages, and that K^+^ efflux operates by enabling FISNA-NACHT conformational flexibility and weakening cage protomer-protomer contacts.

### Physiological K^+^-dependent triggers and autoinflammation-associated mutations render NLRP3 insensitive to potassium stabilization

To test whether the results obtained using manipulating K^+^ concentration in cell lysates, in purified protein solution, or by computational methods also applied to physiological NLRP3 activation under K^+^ efflux conditions in viable cells, we treated cells with nigericin, a classical trigger inducing K^+^ efflux, prior to lysis and pronase digest. First, we used the HEK293T NLRP3 overexpression system and found that pre-treatment of intact cells with nigericin led to an almost complete absence of the NACHT band, even in lysates subsequently prepared in 150 mM K^+^-containing buffer, whilst MCC950 addition to the lysates re-established stabilization in both buffer conditions (Fig. 4A and S5A, quantified in Fig. 4B). This indicated that nigericin treatment of cells, by inducing K+ efflux, reduced the pool of stabilizable NLRP3, an effect that was detectable by limited proteolysis afterwards. To show this in a more physiological setting, we assessed the results of limited proteolysis on endogenous NLRP3 in lysates of LPS-primed THP-1 cells in which the binding of ASC would render opening of the structure less reversible. To assess K^+^ specificity, cells were treated either with nigericin (Fig. 4C, quantified in 4D, and S5B) or with CL097 (Fig. 4E, quantified in 4F, and S5C), an NLRP3 activator that does not trigger K^+^ efflux (Gross, Mishra et al. 2016). A much lower abundance of the NACHT band even in K^+^- or MCC950-containing post-lysis conditions was observed for nigericin compared to CL097 (Fig. 4C/D vs E/F). For the latter, a higher ratio of the protection band was detected, i.e., a greater amount of conformationally compact NLRP3 (cf. Fig. 4D). Hence, complete band disappearance was specific to lysates from cells exposed to the K^+^-dependent NLRP3 trigger nigericin, and not observed in response to a K^+^-independent NLRP3 stimulus. Thus, the conformational transition of isolated NLRP3 (*cf.* Fig. 2A) in pronase assays mirrors what is observed under typical K^+^ efflux conditions of NLRP3 activation in living cells.

**Figure 4:**
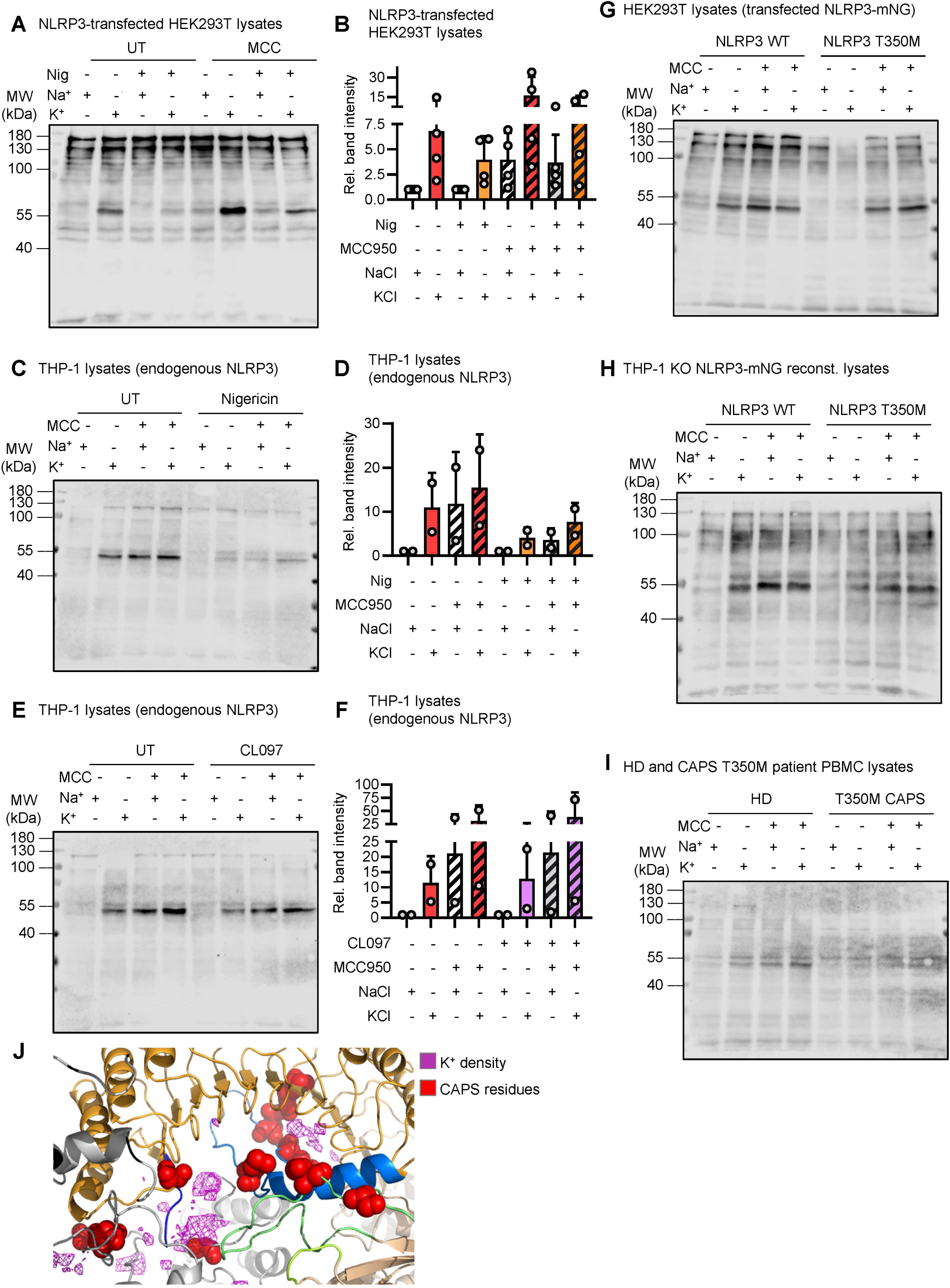
The conformational state of NLRP3 in cells treated with a K^+^ efflux trigger and of disease-associated NLRP3 variants corresponds to the NLRP3 conformation in K^+^-free lysates. **(A)** Immunoblot of NLRP3-mNG-expressing and nigericin-treated HEK293T cell lysates prepared by sonication in K⁺ (150 mM KCl) or Na⁺ (150 mM NaCl) buffer, followed by ± MCC950 and subsequent pronase addition (undigested lysates shown in Fig. S5A). Anti-NLRP3 antibodies were used. Data are representative of n=4 independent experiments. **(B)** Quantification of A combining n=4 independent experiments. **(C)** Immunoblot of LPS-primed, VX-765 pre-treated, and nigericin-treated regular THP-1 cell lysates prepared by sonication in K⁺ (150 mM KCl) or Na⁺ (150 mM NaCl) buffer, followed by ± MCC950 and subsequent pronase addition (undigested lysates shown in Fig. S5B). Anti-NLRP3 antibodies were used. Data are representative of n=2 independent experiments. **(D)** Quantification of C showing combined data from n=2 independent experiments. **(E)** As in C but with CL097 instead of nigericin treatment and addition of 40 mM KCl addition to the media (undigested lysates shown in Fig. S5C). Data are representative of n=2 independent experiments. **(F)** Quantification of E showing combined data from n=2 independent experiments. **(G)** Immunoblot of cell lysates from NLRP3 WT-mNG or NLRP3 T350M-mNG-texpressing HEK293T cells with concomitant MCC950 treatment. Cell lysates were prepared by sonication in K⁺ (150 mM KCl) or Na⁺ (150 mM NaCl) buffer including DMSO or MCC950, followed by pronase addition (quantification in Fig. S5D, undigested lysates shown in Fig. S5E). Anti-NLRP3 antibodies were used. Data are representative of n=3 independent experiments. **(H)** Immunoblot of LPS-primed NLRP3 WT-mNG or NLRP3 T350M-mNG reconstituted THP-1 *NLRP3* KO cell lysates prepared by sonication in K⁺ (150 mM KCl) or Na⁺ (150 mM NaCl) buffer, followed by ± MCC950 for 72 hours and subsequent pronase addition (undigested lysates shown in Fig. S5H). Anti-NLRP3 antibodies were used. Data are representative of n=2 independent experiments. **(I)** Immunoblot of healthy donor or CAPS T350 patient PBMC prepared by sonication in K⁺ (150 mM KCl) or Na⁺ (150 mM NaCl) buffer, followed by ± MCC950 and pronase addition (undigested lysates shown in Fig. S5I). Anti-NLRP3 antibodies were used. n=1 independent experiment with samples from 1 healthy donor and 1 CAPS patient. **(J)** Structure of NLRP3 face-to-face (F2F) dimers extracted from the 7PPZC decameric cage and overlaid with K^+^ computer densities (magenta) calculated from MDS. Selected residues mutated in CAPS are highlighted as red spheres. Data are representative from n=8 independent MDS (*cf*. Fig. 3B).

We sought to relate our findings to NLRP3-associated disease in humans, namely, the autoinflammatory disease CAPS (Weber, McManus et al. 2025). Although in a myeloid cell line (U937 cells) different classes of CAPS mutants show different requirements for hyperactivation (Cosson, Riou et al. 2024), limited proteolysis assay of the known CAPS variants T350M (Fig. 4G, quantified in Fig. S5D, and Fig. S5E) and E569K (Fig. S5F, quantified in S5G) in transfected HEK293T cell lysates showed a strong decrease in the protection band even in K^+^ buffer lysis conditions compared to WT NLRP3, in good agreement with a constitutively hyperactive, K^+^ efflux-independent phenotype of these variants in this not fully inflammasome-competent cell system (Tapia-Abellan, Angosto-Bazarra et al. 2019). However, as previously reported for some of these mutants (Coll, Hill et al. 2019, Tapia-Abellan, Angosto-Bazarra et al. 2019, Weber, Tapia-Abellan et al. 2022), addition of MCC950 concomitant to cell maintenance and transfection, stabilized these variants against proteolytic degradation independently of the lysis buffer used (Fig. 4H and S5H). This was further confirmed using *NLRP3* KO THP-1 cells reconstituted with either WT or T350M CAPS NLRP3-mNG. Interestingly, despite comparable expression, the WT NLRP3-derived protection band was evident in K^+^-containing lysis buffer and after MCC950 pre-treatment, while the T350M variant showed a weak protection band only in the presence of MCC950 (Fig. 4H, quantified in S5H, and S5I). Finally, we sought to confirm this observation in primary immune cells (PBMC) from a CAPS patient heterozygous for the NLRP3 T350M variant. Despite the low sample abundance, we observed a similar trend, namely, that a protection band could only be partially restored by MCC950 addition but not by high K^+^ buffer (Fig. 4I and S5G), in contrast to PBMC from a healthy donor, in which both K^+^ and MCC950 stabilized the unstimulated, conformationally compact state. Interestingly, the proximity of certain CAPS mutants to the nucleotide binding pocket or the acidic loop has been discussed as relevant for auto-activity (Feng, Wierzbowski et al. 2025), and is thus in good agreement with a proximity to MDS-computed K^+^ densities in our model (Fig. 4J). These results are consistent with earlier work (Weber, Tapia-Abellan et al. 2022) and indicate that NLRP3 CAPS variants present a structure that can be stabilized by MCC950 if added co-translationally before lysis to resist degradation, by forcing a compact inactive 3D conformation. However, our data indicate that the presence of a high K^+^ concentration alone appears insufficient to reverse their active and more open conformation, which is thus degraded more easily. Thus, at the conformational level, the difference between closed, inactive WT NLRP3 (unstimulated cells or K^+^-containing lysate) and open, active WT NLRP3 (ASC-competent cells with nigericin stimulation or K^+^-free lysates) mirrors the difference between WT and CAPS NLRP3. Our results link these physiologically and disease-relevant conformational differences to a K^+^ sensing role of NLRP3 and support its destabilization at both the monomeric FISNA-NACHT domain and at the cage level as a plausible mechanistic route that connects K^+^ efflux to inflammasome activation.

## Discussion

The reactivity of NLRP3 to K^+^ efflux (Munoz-Planillo, Kuffa et al. 2013) is one of the most widely held paradigms in the inflammasome field and a common danger sensing mechanism that is almost impossible to underestimate—given the pivotal role of NLRP3 in infection, as well as in acute and chronic inflammation underlying sterile pathologies such as myocardial infarction, stroke, Alzheimer’s or Parkinson’s disease (Weber, McManus et al. 2025); but mechanistically it has remained largely unsolved. This is even more surprising given that the earliest connection with K^+^ was made three decades ago (Perregaux and Gabel 1994), predating the discovery of NLRP3 itself. The direct response of NLRP3 to K^+^ would render NLRP3 the first described intracellular/cytosolic K^+^ danger sensor protein in humans. In general, apart from K^+^ channels, so far only one intracellular K^+^ sensor protein has been described in nature, specifically in prokaryotes, the *E. coli* K^+^ binding protein Kbp (also known as YgaU) (Noskov and Roux 2006, Ashraf, Josts et al. 2016). In *E. coli*, Kbp regulates salt-induced osmotic stress and adopts a compact conformation in the presence of K^+^. Although the structural similarity appears very low, conceptually, the stabilization of inter-domain contacts a common theme shared even with human NLRP3: whereas in the presence of K^+^, the LysM and BON domains of Kbp are held together in a compact, inactive state (Ashraf, Josts et al. 2016), upon K^+^ loss, Kbp structurally ‘unravels’, triggering so far little understood signaling events that prompt a bacterial response to osmotic stress (Bischof, Rehberg et al. 2017). Structurally, loss of K^+^ in NLRP3 seems to cause a more subtle conformational response than in Kbp; nevertheless, critical intra- and intermolecular domain contacts are weakened as a shared mechanism by which K^+^ initiates a cellular response to osmotic (Kpb) or homeostatic stress (NLRP3), respectively.

Our data robustly demonstrate that a functional cell environment – whether in PBMCs, macrophage-like THP-1 cells, NLRP3 reconstituted mammalian HEK293T cells or even non-mammalian, macrophage-like S2 insect cells – is not required for the response of NLRP3 to changes in K^+^ ion concentration. Rather, it appears that K^+^ directly impacts NLRP3 conformation at two levels (**Fig. S6**), FISNA-NACHT and F2F cage interface.

Firstly, at the level of the FISNA-NACHT domain, K^+^ is computed to interact with Thr233, a residue in the ADP/ATP binding pocket which strongly impacts ATP hydrolysis (Brinkschulte, Fussholler et al. 2022). We propose that K^+^ has a stabilizing effect on the ADP-bound structure (including the NBD-HD1/WHD-HD2 interface) similar to NLRP3 small molecule inhibitors that bind in this pocket and thus prevent activation (Dekker, Mattes et al. 2021). Alternatively, K^+^ may affect NACHT stability via Mg^2+^ coordination and subsequent nucleotide coordination. Mechanistically, involvement of the NLRP3 FISNA-NACHT module is in good agreement with domain-swap experiments (Tapia-Abellan, Angosto-Bazarra et al. 2021). We hypothesize that the dissociation of K^+^ from the FISNA-NACHT domain may provide sufficient structural plasticity for nucleotide exchange and the pivoting movement of the NBD-HD1/WHD-HD2 sub-domains.

Apart from effects on individual FISNA-NACHT modules, our analysis also indicates that the F2F interaction within NLRP3 assemblies is stabilized by K^+^, lowering the negative electrostatic field between the protein chains. In line with computational data, removal of K^+^ would thus generate strong electrostatic repulsive force and disrupt this interface (similarly to CAPS variants thought to affect the acidic loop positioning (Feng, Wierzbowski et al. 2025)), whilst the B2B may ensure that the local concentration of NLRP3 remains high enough to form the active disk. We speculate that the compounded weakening effect at the F2F dimer interface – especially the LRR-LRR contacts that need to be resolved for the formation of an active state (Xiao, Magupalli et al. 2023) – may be necessary, or at least conducive, to translate the aforementioned NACHT conformational change into a full ‘breaking open’ of the inactive cage and thus to expose the PYDs for downstream signaling (**Fig. S6**). K⁺ may thus act as a molecular ‘stabilizer’ freezing NLRP3 monomers (via the FISNA-NACHT) and assemblies (via F2F interfaces) in a closed conformation, similar to MCC950 (Hochheiser, Pilsl et al. 2022). Increased FISNA-NACHT and cage plasticity upon K^+^ efflux thus emerges as a new mechanistic framework that seamlessly integrates previous structural studies (e.g. (Andreeva, David et al. 2021, Hochheiser, Pilsl et al. 2022, Xiao, Magupalli et al. 2023, Yu, Matico et al. 2024). Currently, the dynamic study of state transitions, e.g., by NMR combined with stable isotope labelling, particularly with respect to the effects of K⁺, appears out of reach for the MDa NLRP3 cages. However, studies in which K^+^ is included instead of inhibitors to form stable protein species may bring about new valuable insights into the dynamics of NLRP3 protein transition between individual states.

Why K^+^, and not the monovalent Na^+^, selectively prevents the switch to the active conformation is unclear but may be related to the larger size and/or the greater ability of K^+^ for π-electron interactions due to the larger ionic radius. Additionally, the lower hydration energy of K^+^ facilitates desolvation and dynamic coordination within flexible protein environments, thereby enabling rapid and reversible conformational transitions (Dudev and Lim 2014). Although K⁺ appears to have a dominant and sufficient direct effect on NLRP3 in all our assays, we do not dispute that other ions, e.g., chloride (Green, Yu et al. 2018), sodium (Mangan, Akbal et al. 2024), or other environmental factors such as temperature (Karasawa, Komada et al. 2022, Wang, Bertheloot et al. 2025) may have additional indirect or direct effects on NLRP3 conformation and activation as a whole. Moreover, post-translational modifications (reviewed in (O’Keefe, Dubyak et al. 2024) and (Weber, McManus et al. 2025)) may further impact structural plasticity directly, or force NLRP3 into particular environments, e.g., membranes, that impose further external constraints on its 3D structure. Lastly, binding of NEK7 may assist in pushing NLRP3 further along the route to an active disk conformation by stabilizing it (Xiao, Magupalli et al. 2023). Nevertheless, our data indicate that K^+^ efflux appears to be dominant in directing 3D conformation by directly acting upon NLRP3. As much as the numerous described posttranslational and trafficking events, cage assembly at the TGN and subsequent assembly with ASC or caspase-1 at the MTOC (reviewed in (Weber, McManus et al. 2025)), are required for priming, they seem somewhat redundant for the actual and fundamental conformational response of NLRP3 to K⁺, which is evident even in the purified protein itself. It will be interesting to address in future studies to what extent, and by which mechanism, posttranslational modification, trafficking and/or NEK7 binding are essential for ‘translating’ an executed K^+^-dependent conformational switch towards full inflammasome assembly. Our results from lysates of nigericin *vs.* CL097-treated LPS-primed THP-1 cells might cautiously be taken as an indication that there could be at least two conformationally different active states that converge on ASC and caspase-1 recruitment even further downstream. For example, it is conceivable that K^+^-independent stimuli like CL097 may destabilize the NACHT and/or the NLRP3 cage through a direct, redox-mediated effect. It remains to be investigated how this would overcome K^+^-dependent inhibition to reach an open conformation by an alternative conformational route. In this respect, it will be interesting to explore whether the proposed active disk structure determined *ex cellulo* (Xiao, Magupalli et al. 2023) represents one of these two (K^+^- *vs* e.g. redox-induced) more ‘upstream’ open conformations in cells, or rather the ultimate active disk on which both K^+^-dependent and -independent pathways converge. Although our focus was to elucidate the effect of K^+^ loss on the NLRP3 cage-dependent pathway, our mechanism can also accommodate smaller, non-cage assemblies of NLRP3 (Hafner-Bratkovic, Susjan et al. 2018, Mateo-Tórtola, Hochheiser et al. 2023). For example, the conformation of the mostly dimeric Δexon3 NLRP3 was still affected by K^+^ concentration (*cf.* Figs. S2D-H) and responded to classical K^+^ efflux stimuli in cells (Mateo-Tórtola, Hochheiser et al. 2023), indicating that dimer contacts, presumably via a single F2F interface, when combined with the stabilizing effect of K^+^ on the FISNA-NACHT, are sufficiently stable to maintain NLRP3 in an inactive conformation in the presence of K⁺.

How exactly this inhibition is lost in disease-associated NLRP3 CAPS mutations has been unclear, since NLRP3 mutant variants have so far evaded structural determination. While very elegant characterizations of NLRP3 disease-associated alleles have highlighted checkpoints required for them to activate downstream signaling (Cosson, Riou et al. 2024, Feng, Wierzbowski et al. 2025), our proteolysis results in reconstituted systems and patient cells provide the first actual conformational evidence that hyperactive NLRP3 spontaneously adopts an open conformation that seems independent of K^+^. This highlights the critical role of intracellular K^+^, and the ability of WT NLRP3 to bind it, for restricting inflammation. Although NLRP3 inhibition can also be achieved in other ways – e.g. non-active site inhibitors (Hartman, Humphries et al. 2024, Wilhelmsen, Deshpande et al. 2025), molecular glue (Sylvain, Stoehr et al. 2025) or PROTAC (Keuler, Ferber et al. 2025) degradation –, our results square well with the structural properties of inhibitors like MCC950, which seem to ‘freeze’ the naturally K^+^-bound state for effective inhibition of NLRP3-mediated inflammation (Hochheiser, Pilsl et al. 2022). We hope that a more detailed mechanistic understanding (Fig. S6) of its natural activation route may accelerate strategies that restrict NLRP3-limited inflammation to prevent or ameliorate NLRP3-linked pathologic states.

### Limitations of the study

We acknowledge that all the presented experiments were performed in vitro or ex vivo (PBMC), but this is due to the fact that subtle 3D conformations of untagged endogenous NLRP3 to date cannot be probed in living organisms or intact cells. Moreover, highly resolved (<1.5 Å resolution) structural studies on MDa protein oligomers in cells and even isolated are not feasible. However, the similarity of the effects of K^+^ (whether modulated in lysis buffers or by enacting efflux in living cells) on inflammasome-competent, exogenously expressed or endogenous NLRP3 and purified NLRP3, for which 3D information is available (Dekker, Mattes et al. 2021, Hochheiser, Pilsl et al. 2022), suggests that what occurs in cells on the conformational level also applies to purified NLRP3 and vice versa. The MDS analysis shown here, whilst performed using state-of-the-art methods on an advanced microsecond timescale, far exceeds any previous NLRP3 study (Yu, Matico et al. 2024) in terms of computational power, but is inherently only able to recapitulate K^+^ charge and not electron shell size. We also acknowledge that K^+^ coordination sites in NLRP3 cannot be precisely pinpointed due to inherent limitations of site-directed mutagenesis (which only addresses side-chains, whereas K^+^ often interacts with carbonyl groups (Ashraf, Josts et al. 2016)), or the structural methods currently available to study NLRP3. Stoichiometries and affinities of NLRP3 for K^+^ could not be determined as K^+^ cannot be labelled and solution methods are confounded heavily by changes in hydration energies of K^+^, which exceed the expected NLRP3 binding energies.

## Methods

### Chemicals, reagents and buffers

Phorbol 12-myristate 13-acetate (PMA, tlrl-pma), LPS ultrapure from *Escherichia coli* K12 (LPS-EK, tlr-peklps), VX-765 (inh-vx765i-1), nigericin sodium salt (tlr-nig) and CL097 (tlrl-c97) were obtained from InvivoGen. MCC950 (Cat.No. S8930) was purchased from Selleckchem. Hygromycin B (50 mg/mL) was obtained from Invitrogen. The potassium chelator PNaSS (poly(sodium 4-styrenesulfonate)) was acquired from Sigma Aldrich. KCl, RbCl and NaCl from Carl Roth. Pronase (protease, *Streptomyces griseus*) was acquired from Roche.

The buffers used in this study were the following: for THP-1, HEK293T and *Drosophila* S2 cell lines the K^+^ buffer contained 50 mM HEPES pH 7.5, 150 mM KCl, 10 mM MgCl_2_ and 0.5 mM TCEP and the Na^+^ buffer 50 mM HEPES pH 7.5, 150 mM NaCl, 10 mM MgCl_2_ and 0.5 mM TCEP (adapted from (Hochheiser, Pilsl et al. 2022)). For PBMCs and the NLRP3 purified proteins, the K^+^ buffer contained 50 mM HEPES pH 7.5, 150 mM KCl, 1.5 mM MgCl_2_ and 0.5 mM TCEP and the Na^+^ buffer 50 mM HEPES pH 7.5, 150 NaCl mM, 1.5 mM MgCl_2_ and 0.5 TCEP mM. For titration experiments, the concentrations of 50 mM HEPES, 1.5 mM MgCl_2_ and 0.5 mM TCEP were kept constant and the KCl and NaCl mM concentrations as indicated amounting to a total of 150 mM of salt.

### Plasmids and cloning procedures

Human NLRP3 or NLRP6 (Uniprot IDs #Q96P20 and #P59044, respectively) encoding constructs, engineered either with an N-terminal YFP and/or a C-terminal *Renilla* luciferase (Luc) tags, were a kind gift from Pablo Pelegrín, University of Murcia. An ENTRY plasmid containing the human NLRP3 coding sequences was generated by PCR and NLRP3 sequence was also cloned into pENTR20-mNeonGreen (mNG) (kind gift from Kay Oliver Schink, University of Oslo) as described (Mateo-Tórtola, Hochheiser et al. 2023). The NLRP3 constructs expressed in THP-1 cells contain the mNG fluorophore separated by a long flexible linker of 8 amino acids (TSGSGSGSG). Plasmids for expression in *Drosophila* cells were generated by inserting the human NLRP3 coding sequence with N-terminal YFP and C-terminal Luc (*Renilla* luciferase) tags into the pAW vector (DGRC Stock 1127). In brief, a pENTR1A plasmid containing human YFP-NLRP3-Luc was transferred into pAW by LR reaction (Thermo Scientific) following the manufacturer’s instructions. CAPS variants of NLRP3 were introduced by site-directed mutagenesis (QuickChange kit, Agilent) into the above mentioned NLRP3-mNG pENTR20 plasmid. Sequencing was performed to confirm correct amplification and the absence of unwanted mutations. All other plasmids were generated using standard molecular cloning techniques. Detailed cloning procedures are available from the authors.

### Cloning, expression, and purification of full-length and Δexon3 recombinant human NLRP3

Full length, human NLRP3 (3-1036, UniProt accession code Q96P20) WT or mutant with the linker region (exon 3 sequence) deleted (Δ95-134) were essentially purified as described in detail in (Hochheiser, Pilsl et al. 2022). In brief, codon-optimized for *Spodoptera frugiperda*, were cloned in an in-house modified pACE-Bac1 vector containing a N-terminal MBP-tag, followed by a Tobacco etch virus (TEV) protease cleavage site. For recombinant protein expression of the different NLRP3 proteins, 500 mL of *Sf9* insect cells were infected with 3 % v/v viral stock of the second virus passage. Expression cultures were incubated for 48 h at 27°C and 80 rpm, and were subsequently harvested by centrifugation at 2000 rpm for 20 min. Cell pellets were washed with PBS and either subsequently used for protein purification or flash-frozen in liquid nitrogen for storage at −80°C. For protein purification, cell pellets were solubilized in lysis buffer (50 mM HEPES pH 7.5, 150 mM NaCl, 0.5 mM TCEP, 10 mM MgCl_2,_ 1 mM ADP), supplemented with 1 mM phenylmethylsulfonyl-fluoride (PMSF), followed by sonication (10 sec on, 5 sec off for 4 min at 40% intensity) on ice. Cell lysates were centrifuged at 25000 rpm and 4 °C for 1 h and the supernatants were filtered with a 45 µm syringe filter, prior to application onto a lysis-buffer equilibrated 5 mL MBP-trap column (GE Healthcare) attached to an ÄKTA-Start FPLC system. The column was subsequently washed with 10 CVs of lysis buffer, followed by elution of the proteins with 5 CVs of lysis buffer supplemented with 15 mM maltose. Proteins were further purified by size exclusion chromatography (SEC) on a Superose 6 increase 10/300 GL column (GE Healthcare) equilibrated with lysis buffer. Elution fractions were analyzed by Coomassie-stained SDS-PAGE gels.

### Cloning, expression, and purification of FISNA-NACHT human NLRP3

Human monomeric FISNA-NACHT NLRP3 (131–679) was generated as described in detail in (Dekker, Mattes et al. 2021). In brief, the protein was expressed and purified from insect cells (Sf9 cells) with an N-terminal His6- and ZZ-tag separated from the NACHT domain via a TEV-cleavage site. The protein was purified in three steps: a Ni2+-NTA affinity purification, followed by cleavage of the tag with TEV protease, ion exchange and a subsequent size exclusion step. The protein was finally resuspended at 9 mg/mL in 150 mM MES pH 6.5, 10% glycerol, 1 mM TCEP, 1 mM MgCl_2_, 100 µM ADP and 250 mM NaCl.

### Cell lines

HEK293T cells were cultured in complete DMEM (4.5 g l^-1^ glucose, containing 10% heat-inactivated fetal bovine serum (FBS), 2 mM L-Glutamine, 100 U/mL penicillin, 100 µg/mL streptomycin) at 37 °C in 5% CO_2_ and sub-cultured every 2-3 days. HEK293T cells were transfected with the indicated plasmids using Lipofectamine 2000 (Thermo Fisher Scientific) according to the manufacturer’s instructions in the presence of 10 µM MCC950 or 0.1% (v/v) DMSO, as indicated. Twenty-four hours after transfection, cell lysates were either directly prepared in the presence of DMSO or MCC950, or NLRP3 activation was induced prior to lysis by treating transfected HEK293T cells for 2 h in complete DMEM with nigericin at 10 µM. For NLRP3 inhibition, cells were pretreated with MCC950 at 10 µM for 30 min and MCC950 was maintained during the stimulation period. Regular THP-1 cells (kind gift from Thomas Zillinger, Bonn, Germany) were maintained in complete RPMI 1640 (containing 10% FBS, 2 mM L-Glutamine, 100 U/mL penicillin/100 µg/mL streptomycin at 37 °C in 5% CO_2_ and sub-cultured every 3 days. THP-1 Null 2 (InvivoGen) and THP-1 Null 2 *NLRP3* KO (InvivoGen) kind gift from Gloria López-Castejon, Manchester, UK were maintained in complete RPMI 1640 (containing 10% heat-inactivated FBS, 2 mM L-Glutamine, 100 U/mL penicillin,100 µg/mL streptomycin, 100 µg/mL Normocin, and 100 µg/mL of Zeocin) and sub-cultured every 3 days. All THP-1 cell lines used in the present work were first differentiated with 100 ng/mL of PMA overnight at a density of 10*10^6^ in 10 mL in cell culture petri dishes, rested for 2 days and primed with 10 ng/mL of LPS-EK for 4 h. Where necessary, cells were pre-treated for 30 minutes and during stimulation with 2 µg/ml VX-765 and stimulated or not with 10 µM nigericin or 100 µM CL097 with 40 mM extracellular KCl for 4 h. The Schneider 2 (S2) *Drosophila melanogaster* cell line (Schneider 1972) was a kind gift from Dr. Florence Schlotter and Jean-Luc Imler, Institut de Biologie Moléculaire et Cellulaire, Strasbourg. S2 cells were seeded at a cell density of 1.5*10^6^/mL and cultured in suspension without CO_2_ at 27 °C to a cell density of 1.5-3*10^7^/mL in T75 flasks. Routine culture was maintained in Schneider’s Drosophila medium (containing 10% FBS and 100 U/mL penicillin,100 µg/mL streptomycin) and subcultured every 3 days. All cell lines were regularly checked by PCR to avoid *Mycoplasma* contamination.

### Generation of stable WT and T350M NLRP3-mNeonGreen expressing THP-1 cell lines

THP-1 *NLRP3* KO cells were lentivirally transduced with WT or T350M NLRP3-mNG encoding lentiviruses as described in (Mateo-Tórtola, Hochheiser et al. 2023).

### Generation of *Drosophila* Schneider 2 (S2) cells stably expressing human NLRP3

For the generation of stable macrophage-like S2 cell lines with YFP-NLRP3-Luc S2 cells were seeded 1.5*10^6^ /ml in 2 mL cell culture medium in 6-well plates and maintained for 24 h at 27°C. For transfection, 2 µg of YFP-NLRP3-Luc plasmid per well was used, placed in Schneider’s *Drosophila* FBS-free medium and mixed with transfection kit Xtreme HP Gene transfection reagent (Roche) at a ratio of 1:3 and left at RT for 30 min following manufacturer’s instructions. The cells were left to rest for 24 h and then, the transfection media was replaced with 2 mL of fresh cell culture media. After 48 h of resting, selection was started with hygromycin B at 300 µg/mL for 3 weeks until the stable cell culture was established. Two methods were used to validate NLRP3 expression in S2 cells. First, a FACS analysis was performed for the cell lines expressing human YFP-NLRP3-Luc, in which we observed that over 84% of the cells were fluorescent. Furthermore, S2 cell lysates were analyzed by immunoblot to check for the correct expression of human NLRP3 using the mouse monoclonal antibody against *Renilla* luciferase (Sigma MAB4400) at 1:2000.

### Study participants and human blood acquisition

All healthy donors and the CAPS patient included in this study provided their written informed consent before participation. Approval for use of biomaterials was obtained by the local ethics committee of the Medical Faculty of Tübingen in accordance with the principles laid down in the Declaration of Helsinki as well as applicable laws and regulations.

### PBMC isolation and priming

Human peripheral blood mononuclear cells (PBMCs) were isolated from whole blood by density centrifugation with Histopaque-1077 (Sigma Aldrich). Briefly, EDTA-anticoagulated blood samples were layered onto Histopaque-1077 and centrifuged at 540 x *g* for 25 min at room temperature without braking. Afterwards, the PBMC layer was carefully collected, transferred into another tube and washed twice with PBS. PBMCs were then resuspended in complete RPMI-1640 and stimulated with 10 ng/ml of LPS-EK for 4 hours at 37 °C in a 6-well plate with 4 x 10^6^ cells in 2 ml total volume per well.

### Generation of cell lysates for pronase digestion assay

All work was performed on ice unless otherwise stated. THP-1, PBMC, HEK293T and S2 cells were washed with ice-cold PBS. Adherent cells were lifted in 1 mL of ice-cold PBS and carefully detached with the help of a scraper and the cell suspension was transferred into a 15 mL Falcon tube and centrifuged at 400 x *g* at 4 °C for 5 min. The supernatant was discarded and the cells resuspended in 2 mL of ice-cold PBS and centrifuged as described above. Then, the supernatant was discarded, and the cell pellet was resuspended in 2 mL of ice-cold PBS, split in two tubes and centrifuged again at 400 x *g* for 5 min. The supernatant was removed carefully, and 2 mL of the respective ice-cold K^+^ or Na^+^ buffer was added without resuspending the pellet. The cell pellets were centrifuged again at 400 x *g* for 5 min and the supernatant discarded. The volume of the dry pellet was estimated and 5 times that volume (around 300-500 µL) of ice-cold K^+^ or Na^+^ buffer was added and resuspended by gentle pipetting up and down. If cell lysates were made from DMSO- or MCC950-treated cells, the respective cell lysis buffer was supplemented to a final concentration of either 1% (v/v) DMSO or 30-100 µM MCC950. The cell suspension was then placed in a precooled 2 mL tube. The cells were then mechanically disrupted by sonication on ice (Output Control = 4; Duty Cycle = constant; 10 s on and 20 s off) 4 times. The sonicated samples were then centrifuged at 17000 x *g* and 4 °C for 20 min. Finally, the supernatants were transferred into prechilled 1.5 mL tubes and kept at −80 °C until the pronase experiments were performed.

### Pronase limited proteolysis assay

For the pronase assay, protein concentrations were first determined by Bradford assay (Sigma) following manufacturer’s instructions. In total, 25 µg of protein from each cell lysate and condition were resuspended into equal volumes of the respective K^+^ or Na^+^ buffer, and incubated with 30 or 100 µM of MCC950, 0.1% (v/v) DMSO or the indicated concentration of PNaSS for 1 h at 30 °C and 400 rpm in a thermoshaker. Then, the lysates were treated or not with 10 ng of pronase per 1 µg of protein. The mix was vortexed quickly, spun down and incubated for 15 min at RT. Then, the calculated volume of 20x protease inhibitor cocktail (cOmplete mini, EDTA-free, Merck/Roche)l was added to the samples, and these were vortexed again, spun down and incubated for 10 min on ice. Finally, the calculated volumes of 4x LDS sample buffer and 10x Reducing agent (both from Thermo Fisher Scientific) were added and the samples were vortexed and boiled at 95 °C for 5 min (total V_sample_ = 50 µL). The samples were then directly used for immunoblot or stored at −80 °C until use.

### Immunoblot

Immunoblots were performed using Tris-glycine SDS–PAGE gels containing 1% (v/v) 2,2,2-trichloroethanol (TCE, example shown in Fig. S1A, further data available upon request). Before transfer, protein amount per lane was visualized by capturing fluorescence of TCE-protein complexes in a UV transilluminator. The gels were then transferred to nitrocellulose membranes (Bio-Rad) by semi-dry electroblotting. Membranes were blocked with Tris-buffered saline with 0.1 % Tween 20 (TBS-T) + 5 % BSA (blocking buffer) and probed overnight at 4 °C with primary antibody at 1:1000. After three washes with TBS-T for 5 min each, membranes were incubated with HRP-conjugated secondary antibody in blocking buffer for 1 h at room temperature. Then, membranes were washed with TBS-T three times for 10 min each. Proteins were visualized using ECL substrate (ThermoFisher) with chemiluminescent detection (LI-COR Odyssey). The primary and secondary antibodies used for immunoblotting in this study are listed in the table below **Table S1**.

| Antibodies | Name | Use | Company/Catalog number |
| --- | --- | --- | --- |
| α-NLRP3 | NLRP3 (D4D8T) rabbit mAb | 1:1000 | Cell Signaling Technology, 15101 |
| α-NLRC4 | NLRC4 (85-170) rabbit pAb | 1:250 | St. John's Laboratory, STJ29519 |
| α-NLRP6 | NLRP6 rabbit pAb | 1:1000 | Sigma SAB1302240 |
| α-Protein A | Protein A rabbit pAb | 1:50000 | Sigma P3775 |
| α-β actin | β-actin mouse mAb | 1:2000 | Sigma A5441-100UL |
| α-Rabbit IgG-HRP | Goat Anti-Rabbit IgG Antibody (H+L), Peroxidase | 1:5000 or 1:10000 | Vector laboratories Inc./PI-1000 |
| α-Mouse IgG-HRP | Anti-Mouse IgG (H+L), HRP Conjugate | 1:5000 or 1:10000 | Promega/W402B |

If a second protein such as β-actin was detected using the same membrane, the secondary antibodies were deactivated for 15 min at 37 °C with 30% H_2_O_2_. Subsequently, the deactivated membranes were washed three times for 5 min with TBS-T and blocked again in 5% BSA in TBS-T for 1 h at room temperature and incubated with the next selected primary antibody as previously described. Protein bands were quantified with local background correction by using Image Studio software (version 5.2, LicorBio) and expressed relative to the untreated NaCl condition.

### Thermal shift assay

Thermal shift assays were performed as previously described (Sharif, Wang et al. 2019). In brief, recombinant NLRP3 proteins described above were diluted to 0.5 mg/mL in buffers containing 20 mM Tris (pH 7.5), 5 mM MgCl_2_, and various KCl and NaCl concentrations as indicated in the graphs (with a 150 mM total of salt concentration) in presence or absence of 800 µM MCC950 and incubated on ice for 15 min. Recombinant *S. cerevisiae* ULP1 (Addgene #64697, a gift from Hideo Iwai, Institute of Biotechnology, Helsinki, Finland) was purified from *E. coli* and used as a negative control (Guerrero, Ciragan et al. 2015). BL21 (DE3) cells transfected with ULP1 encoding plasmids were grown in LB medium supplemented with 50 µg/kanamycin and protein expression was induced overnight with 0.1 mM isopropyl-β-d-thiogalactopyranoside at 18 °C. Collected cells were resuspended in buffer containing 50 mM Tris, 500 mM NaCl, 5 mM MgCl_2_, 10% glycerol, 20 mM imidazole and 2 mM 2-mercaptoethanol, pH 7.5, and lysed by sonication. ULP1 was purified by affinity chromatography using Ni-NTA beads (Qiagen) followed by a size-exclusion chromatography on a Superdex 200 column (Cytiva) equilibrated with the gel filtration buffer containing 20 mM Tris-HCl, 150 mM NaCl and 2 mM 2-mercaptoethanol, pH 7.5. SYPRO Orange Protein Stain (#S6650, Invitrogen) was added at 1-fold concentration and melting curves were recorded using CFX Connect Real-Time PCR System (Biorad) with a temperature escalation from 25 to 95 °C with an increment of 0.5 °C/10 s. Δ(RFU)/ΔT was plotted and absolute minima recorded as melting temperature (Tm).

### Molecular dynamics simulations

Starting structures of the molecular systems were built based on the cryo-EM structure of the NLRP3 decamer (PDB code 7pzc, resolution 3.9 Å) and the crystal structure of the NLRP3 FISNA-NACHT domain (PDB code 7alv, resolution 2.83 Å). MCC950 was removed, but ADP was maintained in the original structure. Missing loops were added using the graphical interface of YASARA (Krieger and Vriend 2015). During this project we built a variety of molecular systems and performed more than a hundred molecular dynamics simulations (MDS). In order to limit the complexity of the data reported here, we only show the results obtained from MDS of the FISNA-NACHT monomer, NLRP3 dimers (F2F and B2B) and the NLRP3 decamer, which were both built based on PDB entry 7pzc (residues 127-1036). Of particular interest was loop L2 (177-200), which was not resolved in any PDB file of the inactive conformation of NLRP3. However, there is evidence from PDB entry 8ej4 that L2 can form an alpha-helix. Also, during our MDS of the NLRP3 FISNA-NACHT domain (PDB entry 7alv) L2 spontaneously formed an alpha-helix spanning residues 184-190 (data not shown). Therefore, we modelled L2 in the dimer and decamer starting structure as an alpha-helix. Assignment of force field parameters and atom charges and setup of the simulation box were performed using the YASARA GUI for the simulations using the AMBER14 force field (He, Man et al. 2020). For the MDS using the CHARMM36 force field (Lee, Cheng et al. 2016), the CHARMM-GUI was used for setup. In general, the complexes were solvated in 150 mM KCl solution and simulations were performed at 310 K (corresponding to 37 °C) using periodic boundary conditions. In order to optimize computing time, cuboid boxes were used for the simulation of the F2F dimer (160 Å x 100 Å x 100 Å, 165,108 atoms), B2B dimer (130Å x 162Å x 95Å, 202475 atoms) and the decamer (200 Å x 175 Å x 200 Å, 743,222 atoms). To prevent rotation of the solute in the cuboid box, the backbone atoms of residues 800-900 in chain A were restrained to their initial position. A cubic box (212 Å) was used for the decamer systems (892,158 atoms) built with the CHARMM-GUI. Simulations were performed using either YASARA with GPU acceleration in ‘fast mode’ (5 fs time step) (Krieger and Vriend 2015) on ‘standard computing boxes’ equipped e.g. with one 12-core i9 CPU and NVIDIA GeForce GTX 1080 Ti. Simulations using the CHARMM force field (4 fs time step using hydrogen mass repartitioning) were performed using GROMACS (Pronk, Pall et al. 2013) (vers. 2021.1) on the bwHPC (JUSTUS 2) high-performance computing cluster. During the simulations with YASARA, the box size was rescaled dynamically to maintain a water density of 0.996 g/mL. For the simulations using GROMACS, the Parrinello-Rahman pressure coupling scheme was used to keep the pressure at 1 bar. Conformational Analysis Tools (CAT, http://www.md-simulations.de/CAT/) was used for analysis of trajectory data, general data processing, and generation of scientific plots. VMD (Humphrey et al. 1996) was used to generate molecular graphics. Structural files and MDS-generated structures were also analyzed using Pymol (Schroedinger).

### Data analysis, visualizations, statistics and reproducibility

Experimental data were analyzed in Excel 2019, Fiji, GraphPad Prism 8 and Image Studio 5.2. The exact value of n is indicated in the figure legends. Normal distribution in each group was first analyzed using the Shapiro-Wilk test to choose the subsequent parametric test (e.g. ANOVA when comparing multiple groups, or one sample *t*-test for comparisons to control, e.g. Na^+^ only) or non-parametric versions (e.g. Mann-Whitney U when comparing multiple groups, or one sample Wilcoxon signed rank test for comparisons to control, e.g. Na^+^ only). The choice of test is indicated in the figure legends. Data are always presented as mean ± SD, *p*-values (α=0.05) were calculated as indicated in the figure legends using GraphPad Prism and values <0.05 were considered statistically significant. Blots were quantified using ImageStudio software (Li-Cor) and band intensities normalized to Na^+^ only samples. For visual annotation of statistical significance in graphs, the following nomenclature was used: *p < 0.05 and ns, not statistically significant. Schematic figures were generated using Adobe Illustrator, PowerPoint, and/or Biorender.com.

### Resource, materials and data availability

Further information and reasonable requests for resources and reagents should be directed to Alex Weber. All materials generated during this study are included in this article and its supplemental information files. Any material that can be shared will be released via a material transfer agreement for non-commercial usage.

## Acknowledgements

We thank Gloria López-Castejón and Fátima Martin-Sánchez (Institute of Immunology and Inflammation, University of Manchester) for providing THP-1 cells used in this paper, Pablo Pelegrin (University of Murcia) for the NLRP3 and NLRP6 plasmids, Kay Oliver Schink (University of Oslo) for the pENTR20-mNG plasmid, Hideo Iwai (Institute of Biotechnology, Helsinki, Finland) for the *S. cerevisiae* ULP1 expression plasmid, Markus W. Löffler (Institute of Immunology, University of Tübingen) for biobank support and acquiring blood samples, and Libera Lo Presti and Yasemin Kaya for excellent editorial support. The pAW plasmid was acquired from the Drosophila Genomics Resource Center (NIH Grant 2P40OD010949).

## Author contributions

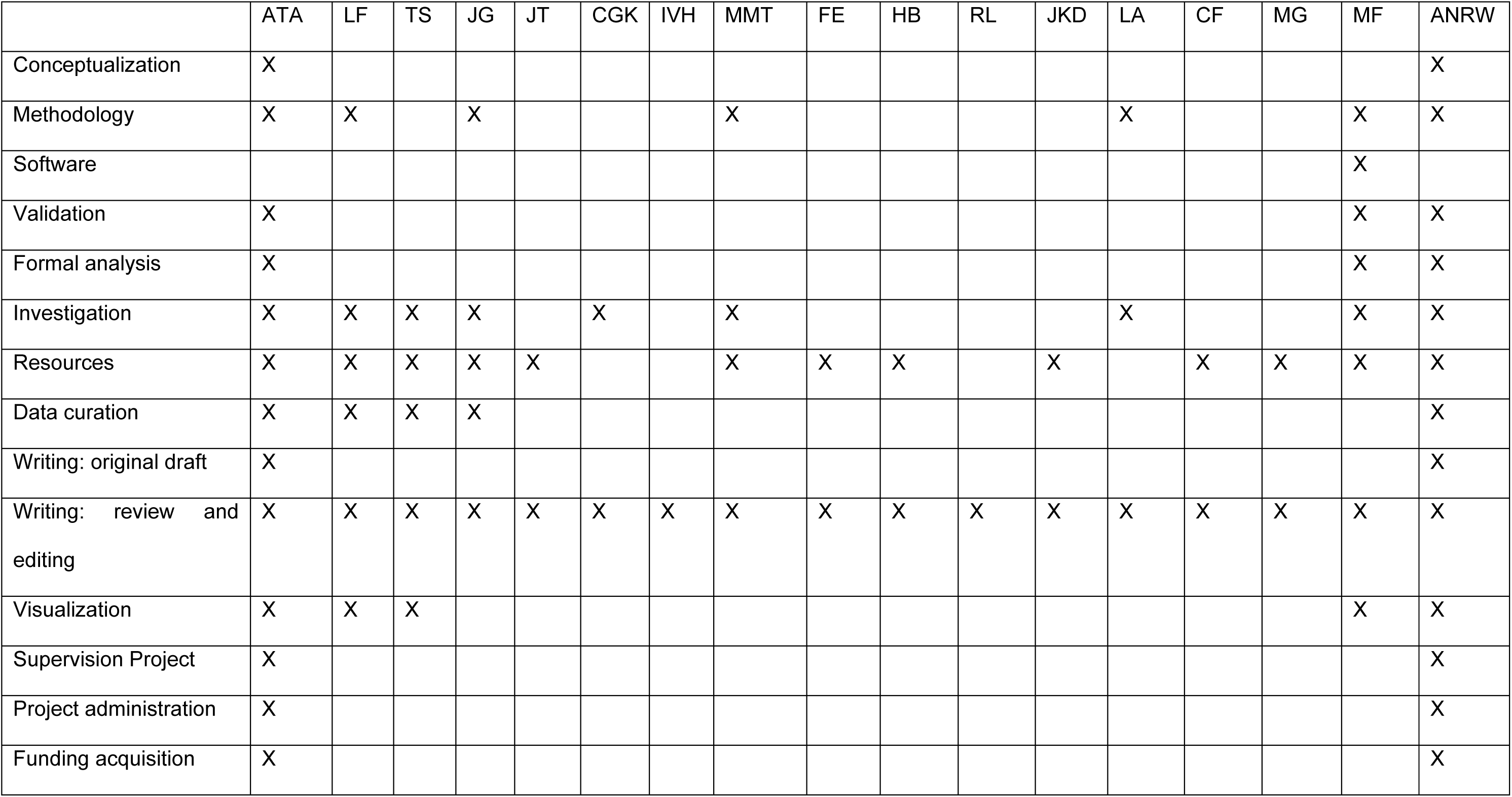

## Supplementary material

**Figure S1: NLRP3 conformation in THP-1 cells and stably transfected *Drosophila* Schneider 2 cells is potassium dependent**

**(A)** 2,2,2-trichloroethanol (TCE) gel analysis of gels shown in Fig. 1A, representative for all gels shown.

**(B)** Immunoblot on LPS-primed regular THP-1 cell lysates prepared by sonication in increasing concentrations of KCl (0 to ∼150 mM) buffer and followed by pronase addition. Anti-NLRP3 antibodies were used. Data are representative of n=3 independent experiments.

**(C)** Quantification of B showing combined data from n=3 independent experiments. One sample t-test.**(D)** As in B but with lysates from *Drosophila* S2 cells stably expressing human YFP-NLRP3-Luc and incubation ± MCC950 or ± PNaSS prior to pronase digestion. Data are representative of n=3 independent experiments.

**(E)** Quantification of D showing combined data from n=3 independent experiments.

**(F)** As in D but using RbCl instead of KCl. Data are representative of n=3 independent experiments.

**(G)** Quantification of F showing combined data from n=3 independent experiments.

**Figure S2: Potassium modulates the conformation of NLRP3 in cell lysates or when recombinantly expressed and purified**

**(A)** Immunoblot of purified recombinant full-length NLRP3 (3-1036) and Protein A in KCl (150 mM) or NaCl (150 mM) buffer, with ± MCC950 added prior to pronase addition (control blot to Fig. 2A). Anti-Protein A antibodies were used. Data are representative of n=5 independent experiments.

**(B)** Quantification of A showing combined data from n=5 independent experiments. One sample Wilcoxon signed rank test.

**(C)** Melting temperature analysis (Tm, °C) of purified recombinant human full-length NLRP3 (3–1036) in buffers containing increasing KCl and decreasing NaCl concentrations (total ion concentration 150 mM) ± MCC950 or ± ADP. Data are representative of n=3 independent experiments.

**(D)** As in A but using anti-NLRP3 antibodies and purified recombinant Δexon3 NLRP3 protein. Data are representative of n=5 independent experiments.

**(E)** Quantification of D showing combined data from n=5 independent experiments. One sample t-test.

**(F)** Immunoblot of lysates from NLRP3 KO THP-1 cells reconstituted stably with Δexon3 NLRP3-mNG. Anti-NLRP3 antibodies were used. Data are representative of n=4 independent experiments.

**(G)** Quantification of F showing combined data from n=4 independent experiments. One sample t-test.

**(H)** As in C but using purified recombinant human Δexon3 NLRP3 protein. Data are representative of n=3 independent experiments.

**(I)** Coomassie stained SDS-PAGE gel of anti-NLRP3 immunoprecipitate from full-length NLRP3-Luc-expressing HEK293T cells subjected to mass spectrometry peptide fingerprinting (Fig. 3F). Data from n=1 independent experiment.

**(J)** Close-up view of the nucleotide binding pocket with two K^+^ ions bound. Selected residues are highlighted.

**Figure S3: Molecular dynamics simulation shows potassium-dependent stability of the F2F interface at dimer and cage level**

**(A)** Trajectory plots showing the stability of ‘native residue-residue contacts’ found in the F2F interface of the NLRP3 face-to-face (F2F) dimers extracted from the 7PZC decameric cage, in the presence (left) and absence (right) of potassium ions. The analysis is based on a two-step procedure where first the native residue-residue contact pairs are determined based on the starting structure. A residue pair is included in the list when any two atoms are closer than a cutoff (5 Å used here). For each residue pair in the list, the distances between the residue centers were calculated and kept as a reference for the analysis. Then, the molecular dynamics trajectories were processed and for each residue pair, the deviation of the distance from the reference distance was calculated. If the current distance deviate**d** less than 4 Å from the reference value, the contact was considered to be ‘native’. Finally, the fraction of native contacts was calculated. A value of 0.7 means that 70% of the residue pairs in the list have distances that are still close to their original values. Combined data from n=8 independent MDS with 0.15 M KCl (letters A-H, left) and n=3 MDS without K^+^ ions (letters A-C, right).

**(B)** As in A but showing the deviation of the residue-residue distances from their reference values as a heat map. Combined data from n=8 independent MDS for 0.15 M KCl (letters A-H, left) and n=3 MDS without K^+^ ions (letters A-C, right).

**(C)** Conformational change observed in MDS ‘A’ (0 mM KCl, see also Fig. 3E, right) resulting in a partially dissociated protein-protein complex. One representative of n=3 identical but independent MDS shown.

**(D)** Simulation of the destabilizing effect of K^+^ efflux on the 7PZC NLRP3 decamer cage interfaces. Trajectories of the distances between residues Pro688 and Tyr1012 in the presence of 0.15 M KCl (left) and in the absence of K^+^ ions (right) extracted from MDS of the 7PZC decameric cage. One representative of n=8 (left) or n=3 (right) identical but independent MDS is shown.

**(E)** As in D (right) but trajectory of the fraction of native residue-residue contacts in the F2F (blue) vs B2B (red) interfaces (average of all five interfaces respectively).

**(F)** Simulation of the destabilizing effect of K^+^ efflux on the 7PZC NLRP3 decamer cage interfaces using GROMACS/CHARMM. Trajectory of the fraction of native residue-residue contacts for the F2F (blue) and B2B (red) interfaces in the presence of 0.15 M KCl (left, average of all five interfaces respectively) or in the absence K^+^ ions (average of all five interfaces respectively)

**Figure S4: Putative involvement of the NLRP3 acidic loop in K^+^ sensitivity**

**(A)** Immunoblot of WT NLRP3-mNG or NLRP3 Δac-loop-mNG-expressing HEK293T cell lysates prepared by sonication in KCl (150 mM) or NaCl (150 mM) buffer, followed by ± MCC950 treatment and subsequent pronase addition for the indicated times (min). Anti-NLRP3 antibodies were used Data are representative of n=1 independent experiments.

**Figure S5: Controls and potassium dependence of NLRP3 CAPS mutants expressed in HEK293T cells**

**(A)** Immunoblot of NLRP3-mNG-expressing and then nigericin-treated HEK293T cell lysates prepared by sonication in K⁺ (150 mM KCl) or Na⁺ (150 mM NaCl) buffer, followed by ± MCC950 addition but without pronase (undigested lysates for Fig. 4A). Anti-NLRP3 antibodies were used. Data are representative of n=4 independent experiments.

**(B)** Immunoblot of LPS-primed, VX-765 pre-treated, and nigericin-treated regular THP-1 cell lysates prepared by sonication in K⁺ (150 mM KCl) or Na⁺ (150 mM NaCl) buffer, followed by ± MCC950 treatment but without pronase addition (undigested lysates for Fig. 4C). Anti-NLRP3 antibodies were used. Data are representative of n=2 independent experiments.

**(C)** As in B but with CL097 instead of nigericin treatment and 40 mM KCl addition to the media (undigested lysates for Fig. 4E). Data are representative of n=2 independent experiments.

**(D)** Quantification of Fig. 4G (Immunoblot of NLRP3 WT-mNG or NLRP3 T350M-mNG--expressing and ± MCC950-treated HEK293T cell lysates) showing combined data from n=3-13 independent experiments. One sample t-test for comparison to Na^+^ only condition; Welch’s t-test for comparison of WT K^+^ with T350M K^+^ condition.

**(E)** Immunoblot of cell lysates from NLRP3 WT-mNG or NLRP3 T350M-mNG-expressing HEK293T cells with concomitant MCC950 treatment. Cell lysates were prepared by sonication in K⁺ (150 mM KCl) or Na⁺ (150 mM NaCl) buffer including DMSO or MCC950 but without pronase addition (undigested lysates for Fig. 4G). Anti-NLRP3 antibodies were used. Data are representative of n=3 independent experiments.

**(F)** As in E but with lysates from NLRP3 WT-mNG- or NLRP3 E569K-mNG-expressing HEK293T cells (undigested and pronase-treated lysates). Data are representative of n=3-13 independent experiments.

**(G)** Quantification of F showing data from n=3-13 independent experiments. One sample Wilcoxon signed rank test.

**(H)** Immunoblot of LPS-primed NLRP3 WT-mNG or NLRP3 T350M-mNG reconstituted THP-1 *NLRP3* KO cell lysates prepared by sonication in K⁺ (150 mM KCl) or Na⁺ (150 mM NaCl) buffer, followed by ± MCC950 for 72 hours but without pronase addition (undigested lysates for Fig. 4H). Anti-NLRP3 antibodies were used. Data are representative of n=2 independent experiments.

**(I)** Immunoblot of PBMC lysates from healthy donor or CAPS T350 patient prepared by sonication in K⁺ (150 mM KCl) or Na⁺ (150 mM NaCl) buffer and followed by ± MCC950 but without pronase addition (undigested lysates for Fig. 4I). Anti-NLRP3 antibodies were used. n=1 independent experiment with samples from 1 healthy donor and 1 CAPS patient.

**Fig. S6: Proposed model of the role of K^+^ in NLRP3 activation.**

Shown are the supposed K^+^-stabilizing effects at the level of the NACHT active site and the F2F decameric cage interface that are lost upon K^+^ efflux, enabling conformational changes in the FISNA-NACHT module and an opening of the cage for assembly of an active inflammasome.

## References

Andreeva, L., L. David, S. Rawson, C. Shen, T. Pasricha, P. Pelegrin and H. Wu (2021). “NLRP3 cages revealed by full-length mouse NLRP3 structure control pathway activation.” Cell 184(26): 6299–6312 e6222.

Ashraf, K. U., I. Josts, K. Mosbahi, S. M. Kelly, O. Byron, B. O. Smith and D. Walker (2016). “The Potassium Binding Protein Kbp Is a Cytoplasmic Potassium Sensor.” Structure 24(5): 741–749.

Bartok, E., M. Kampes and V. Hornung (2016). “Measuring IL-1beta Processing by Bioluminescence Sensors II: The iGLuc System.” Methods Mol Biol 1417: 97–113.

Bischof, H., M. Rehberg, S. Stryeck, K. Artinger, E. Eroglu, M. Waldeck-Weiermair, B. Gottschalk, R. Rost, A. T. Deak, T. Niedrist, N. Vujic, H. Lindermuth, R. Prassl, B. Pelzmann, K. Groschner, D. Kratky, K. Eller, A. R. Rosenkranz, T. Madl, N. Plesnila, W. F. Graier and R. Malli (2017). “Novel genetically encoded fluorescent probes enable real-time detection of potassium in vitro and in vivo.” Nat Commun 8(1): 1422.

Bittner, Z. A., X. Liu, M. Mateo Tortola, A. Tapia-Abellan, S. Shankar, L. Andreeva, M. Mangan, M. Spalinger, H. Kalbacher, P. Duwell, M. Lovotti, K. Bosch, S. Dickhofer, A. Marcu, S. Stevanovic, F. Herster, Y. Cardona Gloria, T. H. Chang, F. Bork, C. L. Greve, M. W. Loffler, O. O. Wolz, N. A. Schilling, J. B. Kummerle-Deschner, S. Wagner, A. Delor, B. Grimbacher, O. Hantschel, M. Scharl, H. Wu, E. Latz and A. N. R. Weber (2021). “BTK operates a phospho-tyrosine switch to regulate NLRP3 inflammasome activity.” J Exp Med 218(11).

Boegli, A., E. M. Bernard, L. Lacante, G. Majeux, E. Hartenian, V. Mack and P. Broz (2026). “The NLRP6 inflammasome is activated by sterile or pathogen-induced endolysosomal damage.” EMBO J 45(1): 30–63.

Boucher, D., M. Monteleone, R. C. Coll, K. W. Chen, C. M. Ross, J. L. Teo, G. A. Gomez, C. L. Holley, D. Bierschenk, K. J. Stacey, A. S. Yap, J. S. Bezbradica and K. Schroder (2018). “Caspase-1 self-cleavage is an intrinsic mechanism to terminate inflammasome activity.” J Exp Med 215(3): 827–840.

Brinkschulte, R., D. M. Fussholler, F. Hoss, J. F. Rodriguez-Alcazar, M. A. Lauterbach, C. C. Kolbe, M. Rauen, S. Ince, C. Herrmann, E. Latz and M. Geyer (2022). “ATP-binding and hydrolysis of human NLRP3.” Commun Biol 5(1): 1176.

Chen, J. and Z. J. Chen (2018). “PtdIns4P on dispersed trans-Golgi network mediates NLRP3 inflammasome activation.” Nature 564(7734): 71–76.

Coll, R. C., J. R. Hill, C. J. Day, A. Zamoshnikova, D. Boucher, N. L. Massey, J. L. Chitty, J. A. Fraser, M. P. Jennings, A. A. B. Robertson and K. Schroder (2019). “MCC950 directly targets the NLRP3 ATP-hydrolysis motif for inflammasome inhibition.” Nat Chem Biol 15(6): 556–559.

Cosson, C., R. Riou, D. Patoli, T. Niu, A. Rey, M. Groslambert, C. De Rosny, E. Chatre, O. Allatif, T. Henry, F. Venet, F. Milhavet, G. Boursier, A. Belot, Y. Jamilloux, E. Merlin, A. Duquesne, G. Grateau, L. Savey, A. T. Jacques Maria, A. Pagnier, S. Poutrel, O. Lambotte, C. Mallebranche, S. Ardois, O. Richer, I. Lemelle, F. Rieux-Laucat, B. Bader-Meunier, Z. Amoura, I. Melki, L. Cuisset, I. Touitou, M. Geyer, S. Georgin-Lavialle and B. F. Py (2024). “Functional diversity of NLRP3 gain-of-function mutants associated with CAPS autoinflammation.” J Exp Med 221(5).

Dekker, C., H. Mattes, M. Wright, A. Boettcher, A. Hinniger, N. Hughes, S. Kapps-Fouthier, J. Eder, P. Erbel, N. Stiefl, A. Mackay and C. J. Farady (2021). “Crystal Structure of NLRP3 NACHT Domain With an Inhibitor Defines Mechanism of Inflammasome Inhibition.” J Mol Biol 433(24): 167309.

Dudev, T. and C. Lim (2014). “Competition among metal ions for protein binding sites: determinants of metal ion selectivity in proteins.” Chem Rev 114(1): 538–556.

Duewell, P., H. Kono, K. J. Rayner, C. M. Sirois, G. Vladimer, F. G. Bauernfeind, G. S. Abela, L. Franchi, G. Nunez, M. Schnurr, T. Espevik, E. Lien, K. A. Fitzgerald, K. L. Rock, K. J. Moore, S. D. Wright, V. Hornung and E. Latz (2010). “NLRP3 inflammasomes are required for atherogenesis and activated by cholesterol crystals.” Nature 464(7293): 1357–1361.

Feng, S., M. C. Wierzbowski, K. Hrovat-Schaale, A. Dumortier, Y. Zhang, M. Zyulina, P. J. Baker, T. Reygaerts, A. Steiner, D. De Nardo, D. L. Narayanan, F. Milhavet, A. Pinzon-Charry, J. I. Arostegui, R. P. Khubchandani, M. Geyer, G. Boursier and S. L. Masters (2025). “Mechanisms of NLRP3 activation and inhibition elucidated by functional analysis of disease-associated variants.” Nat Immunol 26(3): 511–523.

Green, J. P., S. Yu, F. Martin-Sanchez, P. Pelegrin, G. Lopez-Castejon, C. B. Lawrence and D. Brough (2018). “Chloride regulates dynamic NLRP3-dependent ASC oligomerization and inflammasome priming.” Proc Natl Acad Sci U S A 115(40): E9371–E9380.

Gross, C. J., R. Mishra, K. S. Schneider, G. Medard, J. Wettmarshausen, D. C. Dittlein, H. Shi, O. Gorka, P. A. Koenig, S. Fromm, G. Magnani, T. Cikovic, L. Hartjes, J. Smollich, A. A. B. Robertson, M. A. Cooper, M. Schmidt-Supprian, M. Schuster, K. Schroder, P. Broz, C. Traidl-Hoffmann, B. Beutler, B. Kuster, J. Ruland, S. Schneider, F. Perocchi and O. Gross (2016). “K(+) Efflux-Independent NLRP3 Inflammasome Activation by Small Molecules Targeting Mitochondria.” Immunity 45(4): 761–773.

Guerrero, F., A. Ciragan and H. Iwai (2015). “Tandem SUMO fusion vectors for improving soluble protein expression and purification.” Protein Expr Purif 116: 42–49.

Hafner-Bratkovic, I., P. Susjan, D. Lainscek, A. Tapia-Abellan, K. Cerovic, L. Kadunc, D. Angosto-Bazarra, P. Pelegrin and R. Jerala (2018). “NLRP3 lacking the leucine-rich repeat domain can be fully activated via the canonical inflammasome pathway.” Nat Commun 9(1): 5182.

Halle, A., V. Hornung, G. C. Petzold, C. R. Stewart, B. G. Monks, T. Reinheckel, K. A. Fitzgerald, E. Latz, K. J. Moore and D. T. Golenbock (2008). “The NALP3 inflammasome is involved in the innate immune response to amyloid-beta.” Nat Immunol 9(8): 857–865.

Hartman, G., P. Humphries, R. Hughes, A. Ho, R. Montgomery, A. Deshpande, M. Mahanta, S. Tronnes, S. Cowdin, X. He, F. Liu, L. Zhang, C. Liu, D. Dou, J. Li, A. Spasic, R. Coll, M. Marleaux, I. V. Hochheiser, M. Geyer, P. Rubin, K. Fortney and K. Wilhelmsen (2024). “The discovery of novel and potent indazole NLRP3 inhibitors enabled by DNA-encoded library screening.” Bioorg Med Chem Lett 102: 129675.

He, W. T., H. Wan, L. Hu, P. Chen, X. Wang, Z. Huang, Z. H. Yang, C. Q. Zhong and J. Han (2015). “Gasdermin D is an executor of pyroptosis and required for interleukin-1beta secretion.” Cell Res 25(12): 1285–1298.

He, X., V. H. Man, W. Yang, T. S. Lee and J. Wang (2020). “A fast and high-quality charge model for the next generation general AMBER force field.” J Chem Phys 153(11): 114502.

He, Y., M. Y. Zeng, D. Yang, B. Motro and G. Nunez (2016). “NEK7 is an essential mediator of NLRP3 activation downstream of potassium efflux.” Nature 530(7590): 354–357.

Hochheiser, I. V., H. Behrmann, G. Hagelueken, J. F. Rodriguez-Alcazar, A. Kopp, E. Latz, E. Behrmann and M. Geyer (2022). “Directionality of PYD filament growth determined by the transition of NLRP3 nucleation seeds to ASC elongation.” Sci Adv 8(19): eabn7583.

Hochheiser, I. V., M. Pilsl, G. Hagelueken, J. Moecking, M. Marleaux, R. Brinkschulte, E. Latz, C. Engel and M. Geyer (2022). “Structure of the NLRP3 decamer bound to the cytokine release inhibitor CRID3.” Nature 604(7904): 184–189.

Hornung, V., F. Bauernfeind, A. Halle, E. O. Samstad, H. Kono, K. L. Rock, K. A. Fitzgerald and E. Latz (2008). “Silica crystals and aluminum salts activate the NALP3 inflammasome through phagosomal destabilization.” Nat Immunol 9(8): 847–856.

Hwang, I., S. Park, S. Hong, E. H. Kim and J. W. Yu (2012). “Salmonella Promotes ASC Oligomerization-dependent Caspase-1 Activation.” Immune Netw 12(6): 284–290.

Karasawa, T., T. Komada, N. Yamada, E. Aizawa, Y. Mizushina, S. Watanabe, C. Baatarjav, T. Matsumura and M. Takahashi (2022). “Cryo-sensitive aggregation triggers NLRP3 inflammasome assembly in cryopyrin-associated periodic syndrome.” Elife 11.

Katsnelson, M. A., L. G. Rucker, H. M. Russo and G. R. Dubyak (2015). “K+ efflux agonists induce NLRP3 inflammasome activation independently of Ca2+ signaling.” J Immunol 194(8): 3937–3952.

Keuler, T., D. Ferber, J. Engelhardt, C. Steinebach, N. Kirsch, M. Marleaux, G. Weindl, M. Geyer and M. Gutschow (2025). “Degrading the key component of the inflammasome: development of an NLRP3 PROTAC.” Chem Commun (Camb) 61(14): 3001–3004.

Krieger, E. and G. Vriend (2015). “New ways to boost molecular dynamics simulations.” J Comput Chem 36(13): 996–1007.

Lee, J., X. Cheng, J. M. Swails, M. S. Yeom, P. K. Eastman, J. A. Lemkul, S. Wei, J. Buckner, J. C. Jeong, Y. Qi, S. Jo, V. S. Pande, D. A. Case, C. L. Brooks, 3rd, A. D. MacKerell, Jr., J. B. Klauda and W. Im (2016). “CHARMM-GUI Input Generator for NAMD, GROMACS, AMBER, OpenMM, and CHARMM/OpenMM Simulations Using the CHARMM36 Additive Force Field.” J Chem Theory Comput 12(1): 405–413.

Liu, X., T. Pichulik, O. O. Wolz, T. M. Dang, A. Stutz, C. Dillen, M. Delmiro Garcia, H. Kraus, S. Dickhofer, E. Daiber, L. Munzenmayer, S. Wahl, N. Rieber, J. Kummerle-Deschner, A. Yazdi, M. Franz-Wachtel, B. Macek, M. Radsak, S. Vogel, B. Schulte, J. S. Walz, D. Hartl, E. Latz, S. Stilgenbauer, B. Grimbacher, L. Miller, C. Brunner, C. Wolz and A. N. R. Weber (2017). “Human NACHT, LRR, and PYD domain-containing protein 3 (NLRP3) inflammasome activity is regulated by and potentially targetable through Bruton tyrosine kinase.” J Allergy Clin Immunol 140(4): 1054–1067 e1010.

Liu, X., Z. Zhang, J. Ruan, Y. Pan, V. G. Magupalli, H. Wu and J. Lieberman (2016). “Inflammasome-activated gasdermin D causes pyroptosis by forming membrane pores.” Nature 535(7610): 153–158.

Lu, A., V. G. Magupalli, J. Ruan, Q. Yin, M. K. Atianand, M. R. Vos, G. F. Schroder, K. A. Fitzgerald, H. Wu and E. H. Egelman (2014). “Unified polymerization mechanism for the assembly of ASC-dependent inflammasomes.” Cell 156(6): 1193–1206.

Magupalli, V. G., R. Negro, Y. Tian, A. V. Hauenstein, G. Di Caprio, W. Skillern, Q. Deng, P. Orning, H. B. Alam, Z. Maliga, H. Sharif, J. J. Hu, C. L. Evavold, J. C. Kagan, F. I. Schmidt, K. A. Fitzgerald, T. Kirchhausen, Y. Li and H. Wu (2020). “HDAC6 mediates an aggresome-like mechanism for NLRP3 and pyrin inflammasome activation.” Science 369(6510).

Mangan, M. S. J., A. Akbal, S. Rivara, G. Horvath, P. Walch, J. Hegermann, R. Kaiser, F. Duthie, J. Selkrig, R. Gerhard, D. Wachten and E. Latz (2024). “Cytosolic sodium accumulation is a cellular danger signal triggering endocytic dysfunction and NLRP3 inflammasome activation.” bioRxiv: 2024.2003.2026.586774.

Mariathasan, S., D. S. Weiss, K. Newton, J. McBride, K. O’Rourke, M. Roose-Girma, W. P. Lee, Y. Weinrauch, D. M. Monack and V. M. Dixit (2006). “Cryopyrin activates the inflammasome in response to toxins and ATP.” Nature 440(7081): 228–232.

Martin-Sanchez, F., V. Compan and P. Pelegrin (2016). “Measuring NLR Oligomerization III: Detection of NLRP3 Complex by Bioluminescence Resonance Energy Transfer.” Methods Mol Biol 1417: 159–168.

Masters, S. L., A. Dunne, S. L. Subramanian, R. L. Hull, G. M. Tannahill, F. A. Sharp, C. Becker, L. Franchi, E. Yoshihara, Z. Chen, N. Mullooly, L. A. Mielke, J. Harris, R. C. Coll, K. H. Mills, K. H. Mok, P. Newsholme, G. Nunez, J. Yodoi, S. E. Kahn, E. C. Lavelle and L. A. O’Neill (2010). “Activation of the NLRP3 inflammasome by islet amyloid polypeptide provides a mechanism for enhanced IL-1beta in type 2 diabetes.” Nat Immunol 11(10): 897–904.

Mateo-Tórtola, M., I. V. Hochheiser, J. Grga, J. S. Mueller, M. Geyer, A. N. R. Weber and A. Tapia-Abellán (2023). “Non-decameric NLRP3 forms an MTOC-independent inflammasome.” bioRxiv: 2023.2007.2007.548075.

Mayor, A., F. Martinon, T. De Smedt, V. Petrilli and J. Tschopp (2007). “A crucial function of SGT1 and HSP90 in inflammasome activity links mammalian and plant innate immune responses.” Nat Immunol 8(5): 497–503.

Munoz-Planillo, R., P. Kuffa, G. Martinez-Colon, B. L. Smith, T. M. Rajendiran and G. Nunez (2013). “K(+) efflux is the common trigger of NLRP3 inflammasome activation by bacterial toxins and particulate matter.” Immunity 38(6): 1142–1153.

Noskov, S. Y. and B. Roux (2006). “Ion selectivity in potassium channels.” Biophys Chem 124(3): 279–291.

O’Keefe, M. E., G. R. Dubyak and D. W. Abbott (2024). “Post-translational control of NLRP3 inflammasome signaling.” J Biol Chem 300(6): 107386.

Palmer, B. F. and D. J. Clegg (2016). “Physiology and pathophysiology of potassium homeostasis.” Adv Physiol Educ 40(4): 480–490.

Perregaux, D. and C. A. Gabel (1994). “Interleukin-1 beta maturation and release in response to ATP and nigericin. Evidence that potassium depletion mediated by these agents is a necessary and common feature of their activity.” J Biol Chem 269(21): 15195–15203.

Petrilli, V., S. Papin, C. Dostert, A. Mayor, F. Martinon and J. Tschopp (2007). “Activation of the NALP3 inflammasome is triggered by low intracellular potassium concentration.” Cell Death Differ 14(9): 1583–1589.

Pronk, S., S. Pall, R. Schulz, P. Larsson, P. Bjelkmar, R. Apostolov, M. R. Shirts, J. C. Smith, P. M. Kasson, D. van der Spoel, B. Hess and E. Lindahl (2013). “GROMACS 4.5: a high-throughput and highly parallel open source molecular simulation toolkit.” Bioinformatics 29(7): 845–854.

Saller, B. S., S. Wohrle, L. Fischer, C. Dufossez, I. L. Ingerl, S. Kessler, M. Mateo-Tortola, O. Gorka, F. Lange, Y. Cheng, E. Neuwirt, A. Marada, C. Koentges, C. Urban, P. Aktories, P. Reuther, S. Giese, S. Kirschnek, C. Mayer, J. Pilic, H. Falquez-Medina, A. Oelgeklaus, V. G. Deepagan, F. Shojaee, J. A. Zimmermann, D. Weber, Y. H. Tai, A. Crois, K. Ciminski, R. Peyronnet, K. S. Brandenburg, G. Wu, R. Baumeister, T. Heimbucher, M. Rizzi, D. Riedel, M. Helmstadter, J. Buescher, K. Neumann, T. Misgeld, M. Kerschensteiner, P. Walentek, C. Kreutz, U. Maurer, A. S. Rambold, J. E. Vince, F. Edlich, R. Malli, G. Hacker, K. Kierdorf, C. Meisinger, A. Kottgen, S. Jakobs, A. N. R. Weber, M. Schwemmle, C. J. Gross and O. Gross (2025). “Acute suppression of mitochondrial ATP production prevents apoptosis and provides an essential signal for NLRP3 inflammasome activation.” Immunity 58(1): 90–107 e111.

Schmacke, N. A., F. O’Duill, M. M. Gaidt, I. Szymanska, J. M. Kamper, J. L. Schmid-Burgk, S. C. Madler, T. Mackens-Kiani, T. Kozaki, D. Chauhan, D. Nagl, C. A. Stafford, H. Harz, A. L. Frohlich, F. Pinci, F. Ginhoux, R. Beckmann, M. Mann, H. Leonhardt and V. Hornung (2022). “IKKbeta primes inflammasome formation by recruiting NLRP3 to the trans-Golgi network.” Immunity 55(12): 2271–2284 e2277.

Schmid-Burgk, J. L., D. Chauhan, T. Schmidt, T. S. Ebert, J. Reinhardt, E. Endl and V. Hornung (2016). “A Genome-wide CRISPR (Clustered Regularly Interspaced Short Palindromic Repeats) Screen Identifies NEK7 as an Essential Component of NLRP3 Inflammasome Activation.” J Biol Chem 291(1): 103–109.

Schneider, I. (1972). “Cell lines derived from late embryonic stages of Drosophila melanogaster.” J Embryol Exp Morphol 27(2): 353–365.

Sharif, H., L. Wang, W. L. Wang, V. G. Magupalli, L. Andreeva, Q. Qiao, A. V. Hauenstein, Z. Wu, G. Nunez, Y. Mao and H. Wu (2019). “Structural mechanism for NEK7-licensed activation of NLRP3 inflammasome.” Nature 570(7761): 338–343.

Sylvain, A., N. Stoehr, F. Ma, A. Cernijenko, M. Schroder, M. Khoshouei, M. Vogelsanger, M. Schoenboerner, A. Burke, P. Rao, J. M. Solomon, J. Paulk, L. Xu, J. Dawson, D. Begue, P. Lefeuvre, E. Ahrne, A. Hofmann, C. J. Dickson, P. Arabin, A. Zimmerlin, M. Kiffe, M. Froehlicher, T. Boersig, A. Elhajouji, M. Brichet, S. Menon, S. Liu, M. Mueller, C. A. Wartchow, J. Lin, Y. C. Gloria, S. Dickhofer, A. N. R. Weber, T. Welzel, J. Kuemmerle-Deschner, C. J. Farady, R. Pulz, F. Bornancin, D. L. Buckley and Z. I. Bassi (2025). “A cereblon-based glue degrader of NEK7 regulates NLRP3 inflammasome in a context-dependent manner.” Cell Chem Biol 32(7): 955–968 e913.

Tapia-Abellan, A., D. Angosto-Bazarra, C. Alarcon-Vila, M. C. Banos, I. Hafner-Bratkovic, B. Oliva and P. Pelegrin (2021). “Sensing low intracellular potassium by NLRP3 results in a stable open structure that promotes inflammasome activation.” Sci Adv 7(38): eabf4468.

Tapia-Abellan, A., D. Angosto-Bazarra, H. Martinez-Banaclocha, C. de Torre-Minguela, J. P. Ceron-Carrasco, H. Perez-Sanchez, J. I. Arostegui and P. Pelegrin (2019). “MCC950 closes the active conformation of NLRP3 to an inactive state.” Nat Chem Biol 15(6): 560–564.

Wang, W., D. Bertheloot, J. Zhang, M. A. Munoz, S. Xu, A. Stennett, C. M. McKee, R. Knight, E. C. McKay, B. S. Franklin, M. J. Rogers, A. K. Bronowska and R. C. Coll (2025). “NLRP3 is a thermosensor that is negatively regulated by high temperature.” bioRxiv: 2025.2009.2029.679254.

Weber, A. N. R., R. M. McManus, V. Hornung, M. Geyer, J. B. Kuemmerle-Deschner and E. Latz (2025). “The expanding role of the NLRP3 inflammasome from periodic fevers to therapeutic targets.” Nat Immunol 26(9): 1453–1466.

Weber, A. N. R., A. Tapia-Abellan, X. Liu, S. Dickhofer, J. I. Arostegui, P. Pelegrin, T. Welzel and J. B. Kuemmerle-Deschner (2022). “Effective ex vivo inhibition of cryopyrin-associated periodic syndrome (CAPS)-associated mutant NLRP3 inflammasome by MCC950/CRID3.” Rheumatology (Oxford) 61(10): e299–e313.

Wilhelmsen, K., A. Deshpande, S. Tronnes, M. Mahanta, M. Banicki, M. Cochran, S. Cowdin, K. Fortney, G. Hartman, R. E. Hughes, R. Montgomery, C. P. Portillo, P. Rubin, T. Salazar, Y. Wang, S. Yan, B. A. Morgan, A. Duisembekova, R. Riou, M. Marleaux, I. V. Hochheiser, H. Buthmann, D. Ferber, J. Torp, W. Wang, M. Cranston, C. M. McKee, T. J. Mawhinney, E. C. McKay, F. K. Eroglu, J. Kummerle-Deschner, A. N. R. Weber, B. F. Py, M. Geyer and R. C. Coll (2025). “Discovery of potent and selective inhibitors of human NLRP3 with a novel mechanism of action.” J Exp Med 222(11).

Xiao, L., V. G. Magupalli and H. Wu (2023). “Cryo-EM structures of the active NLRP3 inflammasome disc.” Nature 613(7944): 595–600.

Yu, X., R. E. Matico, R. Miller, D. Chauhan, B. Van Schoubroeck, K. Grauwen, J. Suarez, B. Pietrak, N. Haloi, Y. Yin, G. J. Tresadern, L. Perez-Benito, E. Lindahl, A. Bottelbergs, D. Oehlrich, N. Van Opdenbosch and S. Sharma (2024). “Structural basis for the oligomerization-facilitated NLRP3 activation.” Nat Commun 15(1): 1164.

Zhao, Y., J. Yang, J. Shi, Y. N. Gong, Q. Lu, H. Xu, L. Liu and F. Shao (2011). “The NLRC4 inflammasome receptors for bacterial flagellin and type III secretion apparatus.” Nature 477(7366): 596–600.

