## Supplementary figures and images for "NLRP3 acts as a direct sensor of intracellular potassium ions"

### Figure S1

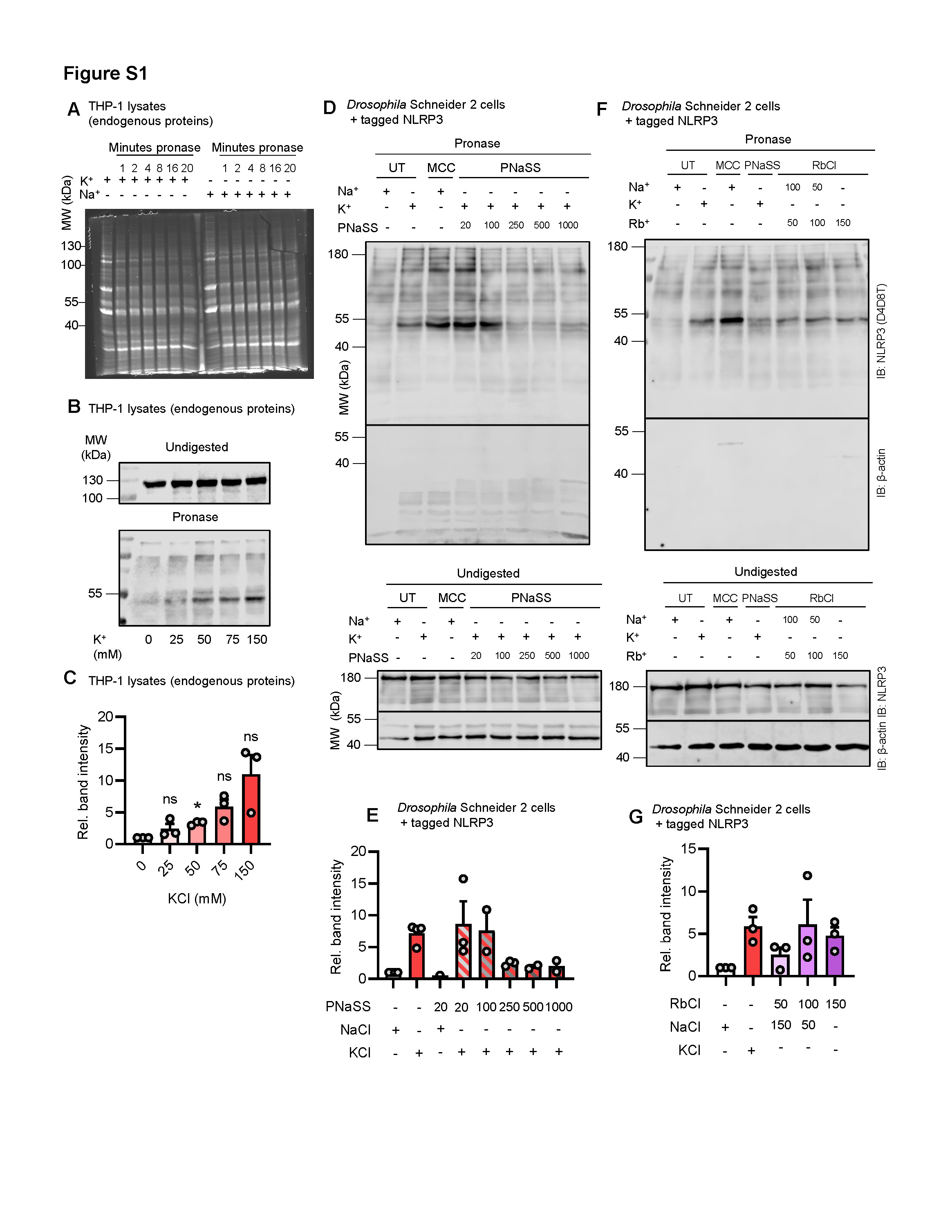

### Figure S2

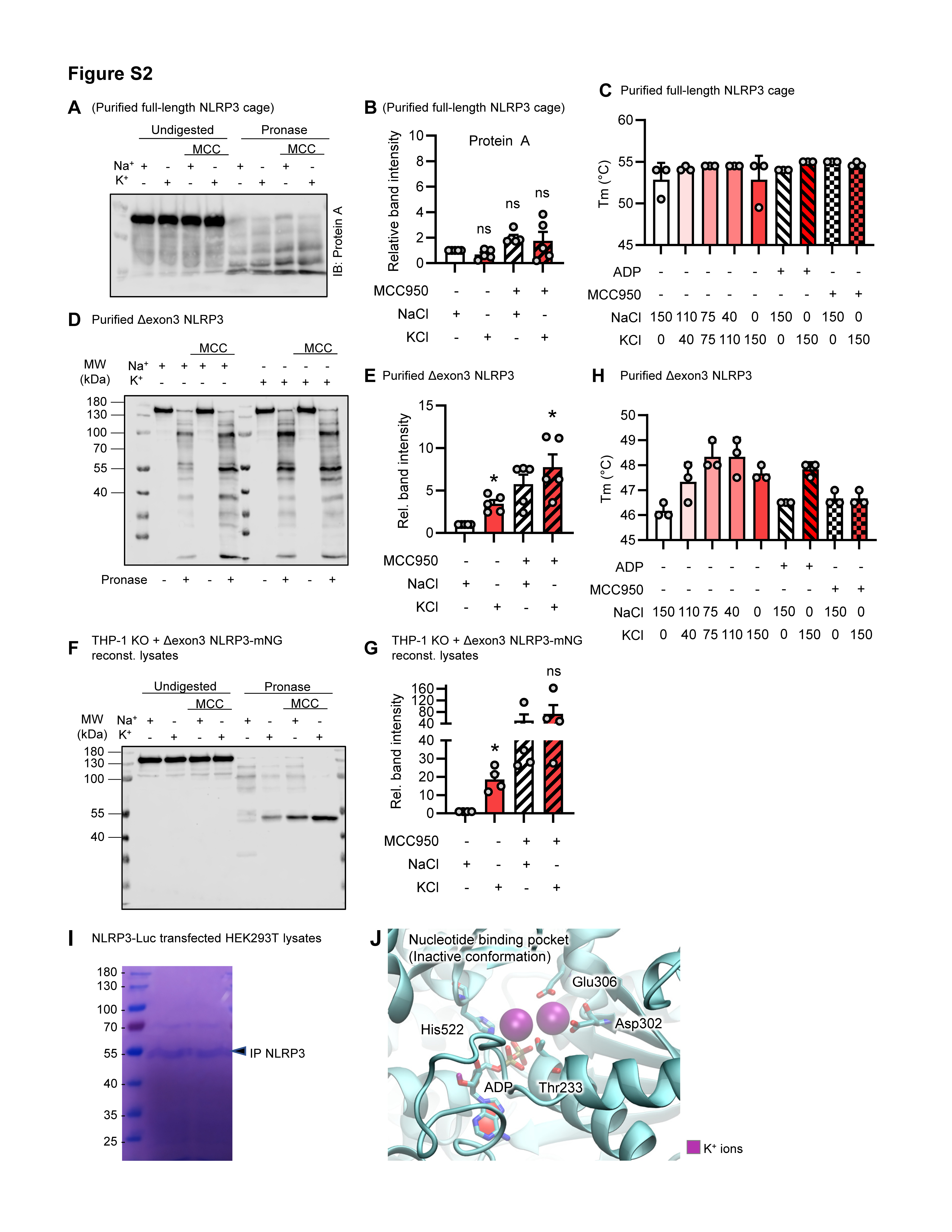

### Figure S3

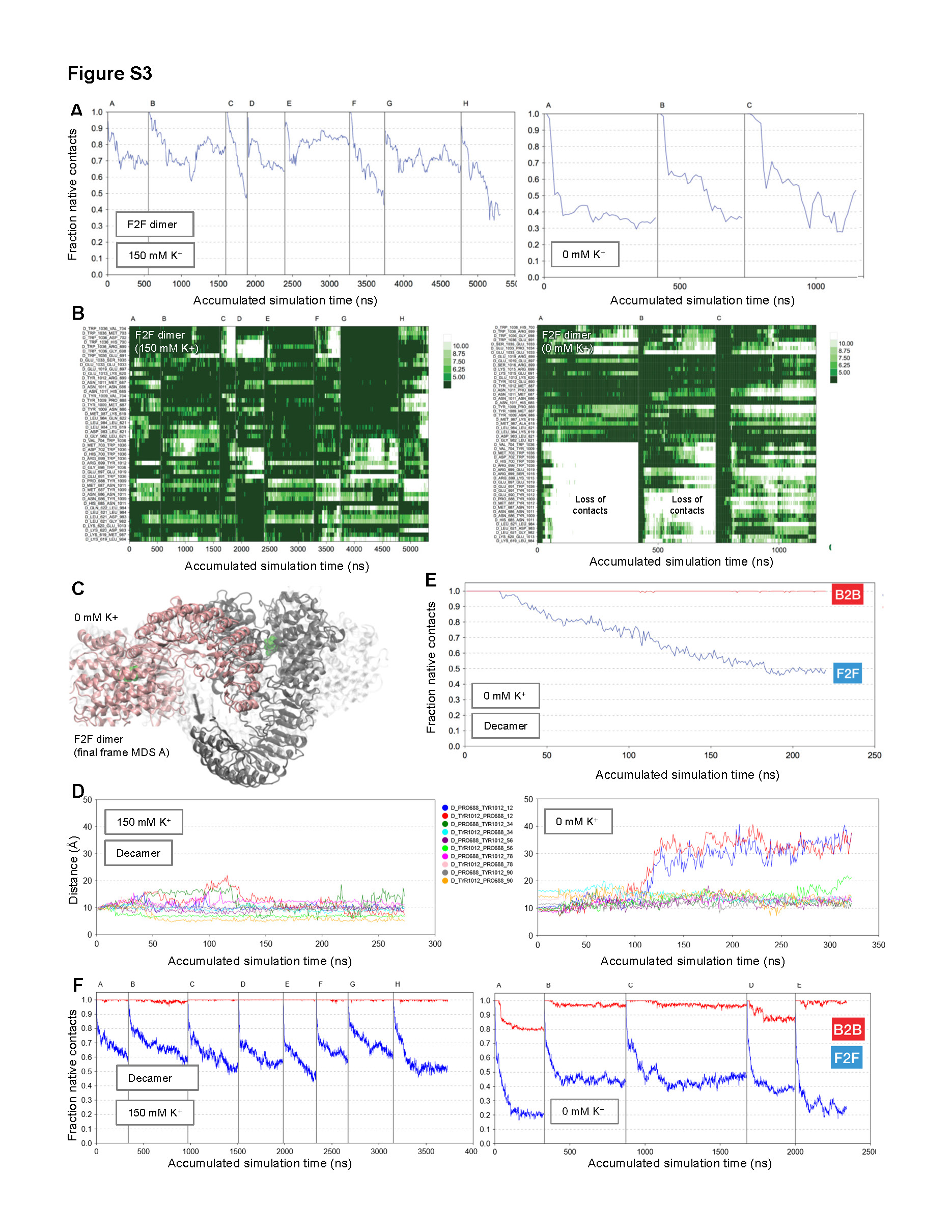

### Figure S4

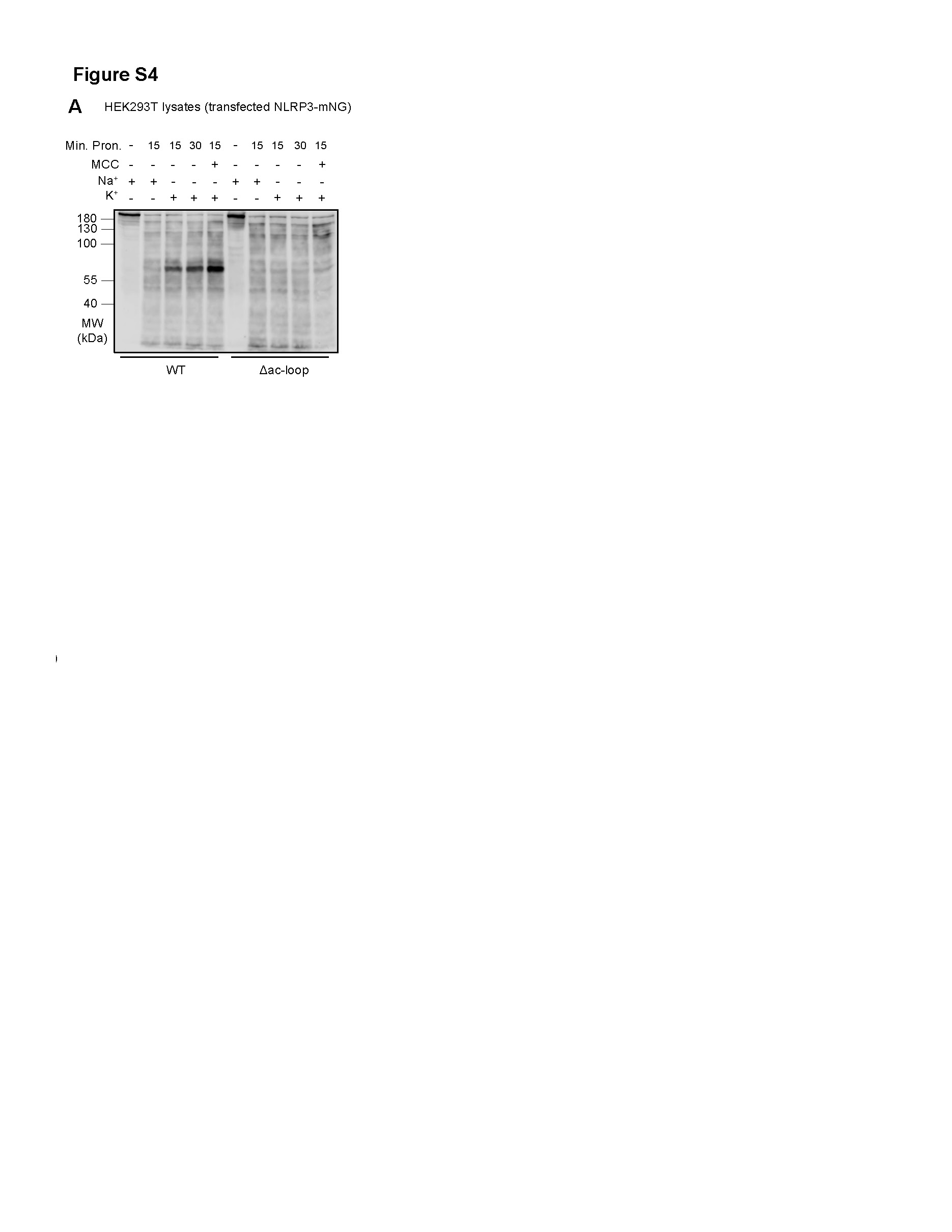

### Figure S5

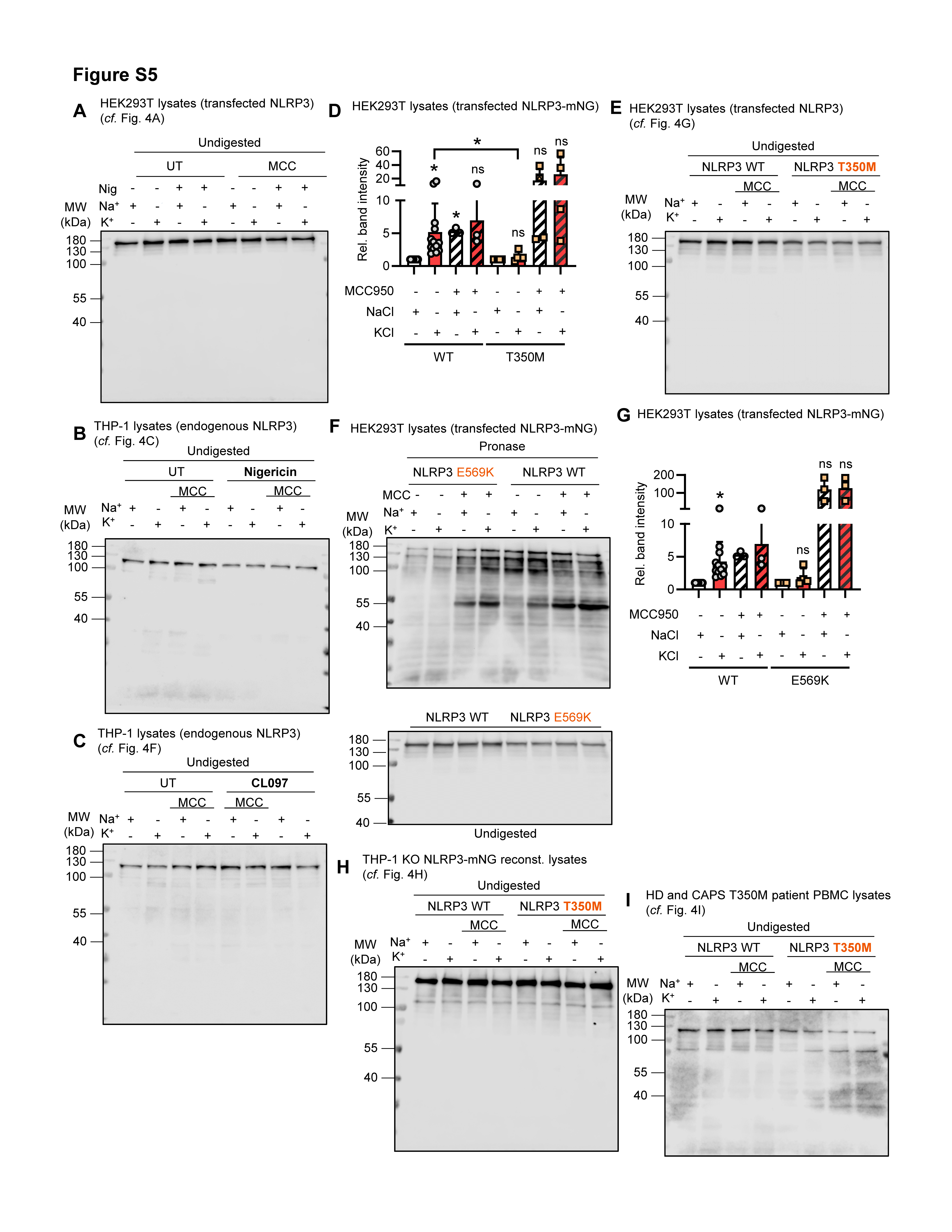

### Figure S6

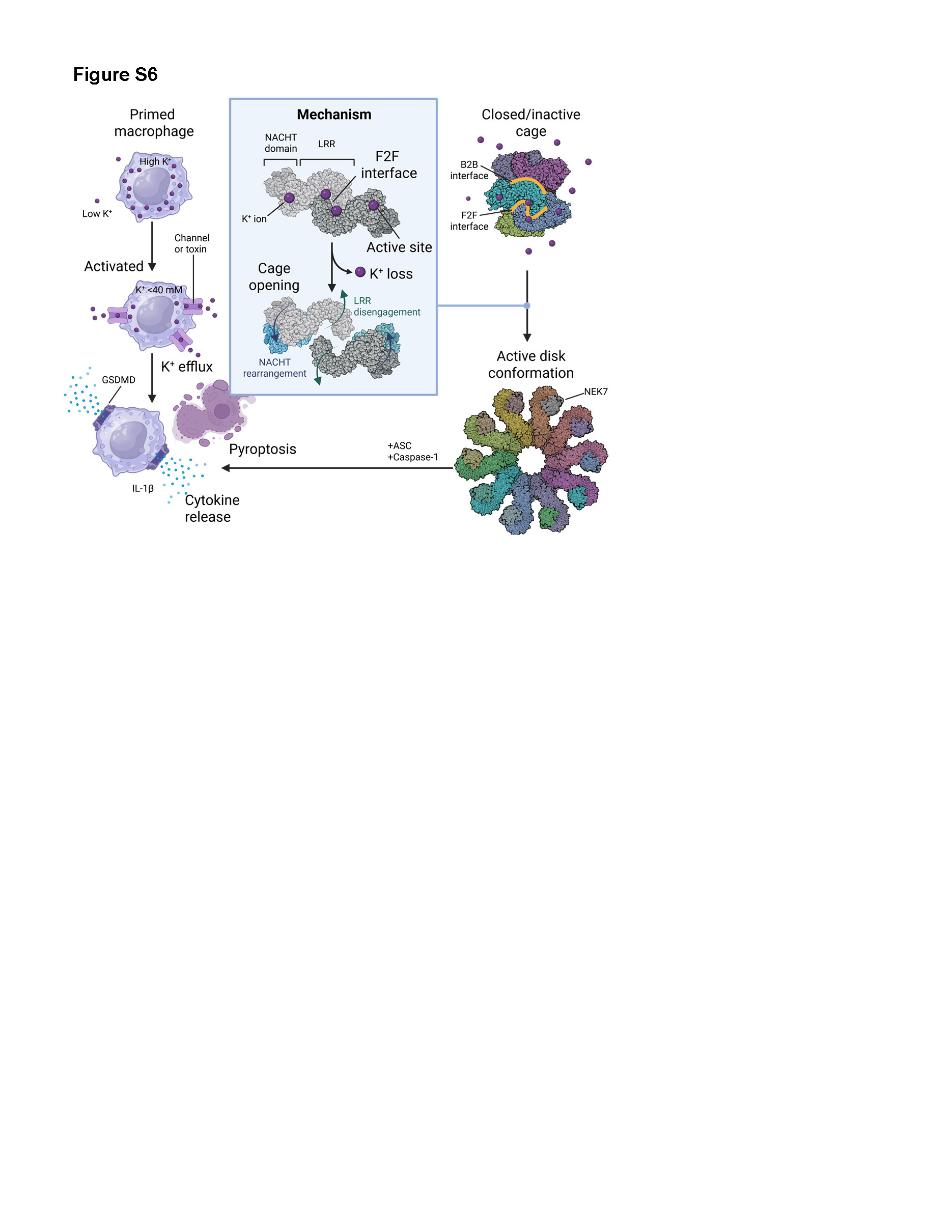
